# KiF1a-regulated neuronal infrastructures in sensory and prefrontal cortices essential for fear-relevant anxiety

**DOI:** 10.1101/2025.07.29.667346

**Authors:** Jiayi Li, Bingchen Chen, Yun Zhang, Yang Xu, Lei Wang, Jin-Hui Wang

## Abstract

Stress in social activities induces fears. Cellular infrastructures and molecular profiles underlying fear-relevant psychosis remain elusive. By giving psychological stress to mice that watched fear scenes in the resident/intruder paradigm, we have studied the features of neuronal infrastructures in visual, auditory and medial prefrontal cortices correlated to fear-induced anxiety by approaches of behavior task, neural tracing, molecular biology and electrophysiology *in vivo*. This psychological stress causes observational mice fears specific to resident mouse and anxiety. This social phobia is associated with the interconnections newly emerged among visual, auditory and medial prefrontal cortices as well as among intramodal neurons. These cortical neurons receive the new synapse innervations from their interconnected neurons alongside the innate synapse innervations, and become to encode the stress signals including battle image and battle sound. The *KiF1a* knockdown in the medial prefrontal cortex precludes the formations of neuronal infrastructures and social phobia. KiF1a-mediated intracellular transportation in the medial prefrontal cortex is essential for the recruitment of associative memory neurons that encode fear scenes and anxiety.

**Highlight:**

1. Psychological stress induces fears and interconnections among sensory and prefrontal cortices.
2. Associative memory neurons in these cortices are recruited to encode fear scenes and anxiety.
3. KiF1a-mediated transportation essential for these cellular infrastructures relevant to learned fear.

## Introduction

The stress in social activities leads to fear memories and phobia. This acquired fear may endorse adaptative functions by activating defensive behaviors in the anticipation of dangerous outcomes and minimizing the impact of threats ^1–3^. This acquired fear by severe persistent stresses also triggers posttraumatic stress disorder, obsessive-compulsive disorder and generalized anxiety ^4–12^. These affective disorders in turn suppress the immune and cardiovascular systems to cause secondary diseases ^13^. It is crucial to reveal cellular and molecular mechanisms underlying stress-induced fears and their impacts, which helps to develop the strategies for the maintenance of moderate fear memory and the relief of stress-induced pathological mood ^1,14–20^.

The acquired fears may occur in the physical stress by body impairment and/or the psychological stress by observing and/or listening fear episodes ^1–3,21^. The neuronal correlates for the acquired fear memory have been studied by using the fear conditioning for rodents to physically receive electrical shocks to their feet with the bell ring or contexture ^8,22–25^. The amygdala, the nucleus accumbens and the medial prefrontal cortex are thought of as critical areas in relevance to the acquired fear ^1,26–55^.

The acquired fear based on body impairment is also induced in the social stress by the resident/intruder paradigm ^7,21,56–63^. Stressful signals including body injury and battle sound are detected by sensory receptors, inputted by afferent nerves, encoded in sensory cortices as well as integrated in association cortices, medial prefrontal cortex and amygdala ^21,64,65^. Those studies through gene screening have shown activity-dependent genes, neuron-building molecules and intracellular signaling cascades relevant to the fear induced by body impairment ^7,60–62,66–70^. As body injury is associated with pain, the molecular and cellular mechanisms underlying pain and pain-relevant fear are overlapped in these data, neither purely for pain nor fear ^18,71^.

In comparison with the pain-relevant fear, the acquired fear more commonly occurs in observing and listening the stress episodes that cause psychological stress ^1–3,72–74^. The neural locations and molecular profiles for the fear induced by psychological stress have been studied ^2,7,60,61,75–78^. In addition to the amygdala ^3,74^, the auditory, visual and medial prefrontal cortices are involved in encoding the stressful signals and fear memory ^21,42,65,79–85^. It remains unclear what are the neuron infrastructure within local cortical areas and the interconnection among cerebral cortices essential for the fear induced by the psychological stress, how the memory traces are constructed to store the psychological fear in the brain, and how the cortical neurons are recruited to encode stress signals and fear-relevant psychosis, which are expectedly addressed in the present study.

The strategies to address such issues are given below. A C57 mouse as the observational subject (observer) was placed in face to a cage of resident/intruder paradigm, in which a resident CD1 mouse attacked an intruder C57 mouse, to induce the fear by the psychological stress ^7,56–59,63,86^. The formation of the acquired fear was assessed by social interaction test and elevated-plus maze test. The interconnections among visual, auditory and medial prefrontal cortices were examined by the neural tracing with adeno-associated virus (AAV) vector that carried the genes of encoding fluorescent proteins. Cellular infrastructures within these cortices were investigated by the neural tracing to track axon interconnections and synapse formations among such cortical neurons and by the *in vivo* electrophysiological recordings in these cortical neurons to analyze their encoding of the stressful signals including sounds and images in battle scenes. The recruitment of associative memory neurons was ensured when new and innate synapse contacts were detected convergently onto dendritic spines of axon-targeted neurons ^21,87–89^ and when these cortical neurons became responding to the fear signals ^87,90,91^. The essential role of the associative memory neurons whose recruitment required motor protein kinesin for intracellular transportation in the acquired fear was examined by using RNAi specific to silence kinesin family member-1a (*KiF1a*).

## Results

The data about cellular infrastructure in medial prefrontal, visual and auditory cortices related to the fear induced by psychological stress is presented in this section. The fear in C57 juvenile mice was onset after witnessing the attacks of a resident mouse to an intruder mouse in the resident/intruder paradigm, in which the stressful signals included scare images and sounds in their battle. The synapse interconnection was morphologically studied by neural tracing based on AAV vectors. The recruitment of associative memory neurons was ensured by both *in vivo* electrophysiological recording and neural tracing, when the cortical neurons receive convergent synapse innervations from the axon boutons of new and innate pathways as well as became to encode stressful signals. The essential role of KiF1a in these processes was examined by shRNA specific for *KiF1a* knockdown.

### Psychological stress by witnessing resident/intruder paradigm induces fear-relevant anxiety

In face to the cage of resident/intruder paradigm once a day for twelve days (a timeline in Figure 1A), observational mice (observers) were assessed by the social interaction test to examine the onset of fear memory specific to this resident mouse and by the elevated-plus maze test to examine anxiety. Compared to control mice (left panel in Figure 1B), observers appear to stay away from this resident mouse in a small interaction box (right panel). The stay durations in an interaction zone, or interaction time (seconds), in response to this resident mouse out of the total time (360 seconds) are 124.5 ±8.09 in observation mice (n=35) and 159.8±9 in controls (n=30, p=0.005, ANOVA, Figure 1C). In comparison with control mice (left panel in Figure 1D), those observers appear to stay away from the battle sound (right panel). The interaction time (seconds) in response to battle sounds is 100.3±5.25 in observers (n=36) and 123.9±6.6 in controls (n=27, p=0.007, ANOVA, Figure 1E). It is noteworthy that the control and observational mice appear not fear to resident mouse and battle sounds before experiments, and that the observers appear not fear to mouse surrogates and sound in their living environment (Figure S1). These results indicate that the mice of experiencing psychological stress express fear memories specific to the resident mouse and battle sounds. Furthermore, in comparison with control mice (left panel in Figure 1F), observational mice appear to avoid the open fields since they prefer to stay in the closed arms of the elevated-plus maze, EPM (right panel). The percentages of the stay time in open arms over the total time on the EPM are 1.76±0.5% in observers (n= 16) and 7.87±1.6 % in controls (n=15, p=0.0008, ANOVA, Figure 1G), indicating that observational mice become more anxious. Thus, psychological stress leads to fear memory and anxiety-like behaviors. As fear memory emerges ahead of anxiety in experiencing a psychological stress (Figure 1H), the anxiety is presumably induced by the fear (Figure 1I).

**Figure 1.**
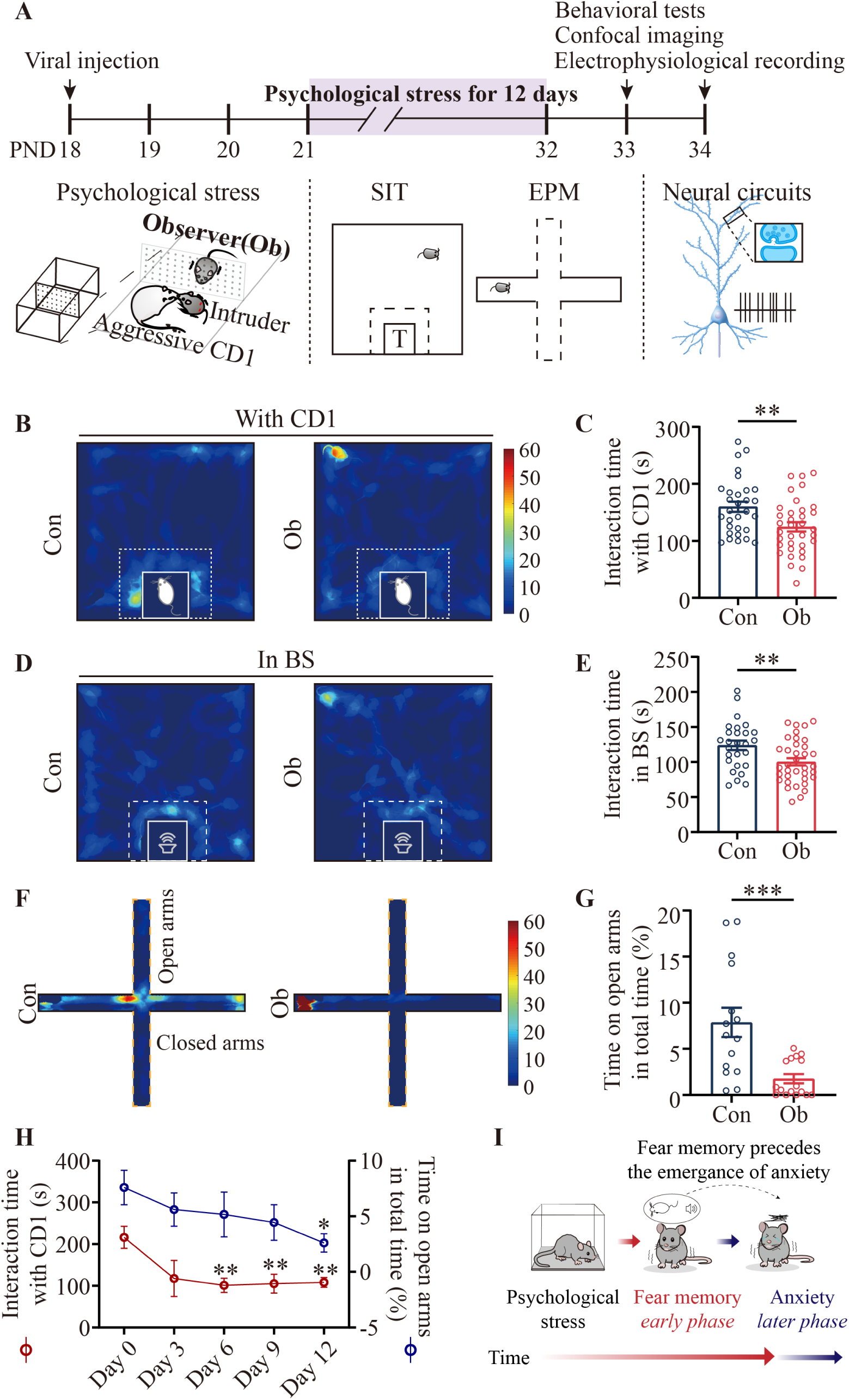
Psychological stress induces fear memory and anxiety-like behavior in mice. **A)** Experiment timeline and diagrams for mice to witness resident/intruder paradigm and to be examined by social interaction test, elevated-plus maze test and cellular plasticity. **B)** Heat maps illustrate the social interaction test for C57 control (Con) and observational (Ob) mice in face to a CD1 resident mouse. **C)** The bar graph with samples’ dots shows the interaction time of C57 mice with CD1 resident. **D)** Heat maps illustrate the social interaction test for C57 mice in response to battle sound. **E)** A bar graph with samples’ dots shows the interaction time of C57 mice with the battle sound. **F)** Heat maps illustrate the elevated-plus maze test for control and observational mice. **G)** A bar graph with sample dots shows the time spent on open arms in the total time on the elevated-plus maze. **H)** A plot shows the social interaction time (left y-axis, red symbols) and the time spent on the open arms in the EPM (right y-axis, blue symbols) measured on psychological stress days 0, 3, 6, 9, and 12. Statistical comparisons were conducted based on the data from time points versus the data on day 0. **I)** A scheme shows that psychological stress induces fear memory and anxiety sequentially. Abbreviations: PND, postnatal day; EPM, elevated plus maze; Ob, observer; Con, control; BS, battle sound. Data are presented as mean ± SEM and analyzed by one-Way ANOVA. Three asterisks, p<0.001; two asterisks, p<0.01.

In terms of cellular infrastructures underlying acquired fears in observational mice, we assumed that the auditory cortex was activated by battle sounds and the visual cortex was activated by battle scenes during the resident/intruder paradigm simultaneously. Based on the rule of coactivity together and interconnection together that recruits associative memory cells ^19,91,92^, we further hypothesized that the stress signals instigated the new synapse interconnections among auditory, visual and prefrontal cortices to recruit these cortical neurons to be associative memory neurons that specifically encode stressful signals, fears and anxiety.

### Psychological stress recruits new synapses and associative memory neurons to encode fear signals

The synapse interconnections between auditory and visual cortices and the synapse innervations from such cortices toward the medial prefrontal cortex were examined by the neural tracing with AAV vector. 0.3 µl AAV-CMV-tdTomato was injected in the visual cortex as well as 0.3 µl AAV-CMV-BFP was injected in the auditory cortex (Figure 2A). Three days after AAV injections, these mice observed the attack of CD1 resident mouse to C57 intruder mouse in resident/intruder paradigm to experience the psychological stress or stayed in a witness cage without C57 intruder mouse in controls once a day for twelve days. The brain slices including visual, auditory and medial prefrontal cortices from two groups were scanned under the confocal microscope. In the visual cortex, the contacts of BFP-labelled axon boutons on YFP-labelled dendritic spines were thought of the synapses made by the axons projected crossmodally from the auditory cortex, and the contacts of RFP-labelled axon boutons on YFP-labelled dendritic spines were the synapses made by the axons among intramodal visual cortical neurons. In the auditory cortex, the contacts of RFP-labelled axonal boutons on YFP-labelled dendritic spines were the synapses made by the axons crossmodally from the visual cortex, and the contacts of BFP-labelled axon boutons on YFP-labelled dendritic spines were the intramodal synapses. In the medial prefrontal cortex, the contacts of BFP- and RFP-labelled axons on YFP-labelled dendritic spines were convergent synapse innervations made by the axons crossmodally from the auditory cortex and the visual cortex.

**Figure 2.**
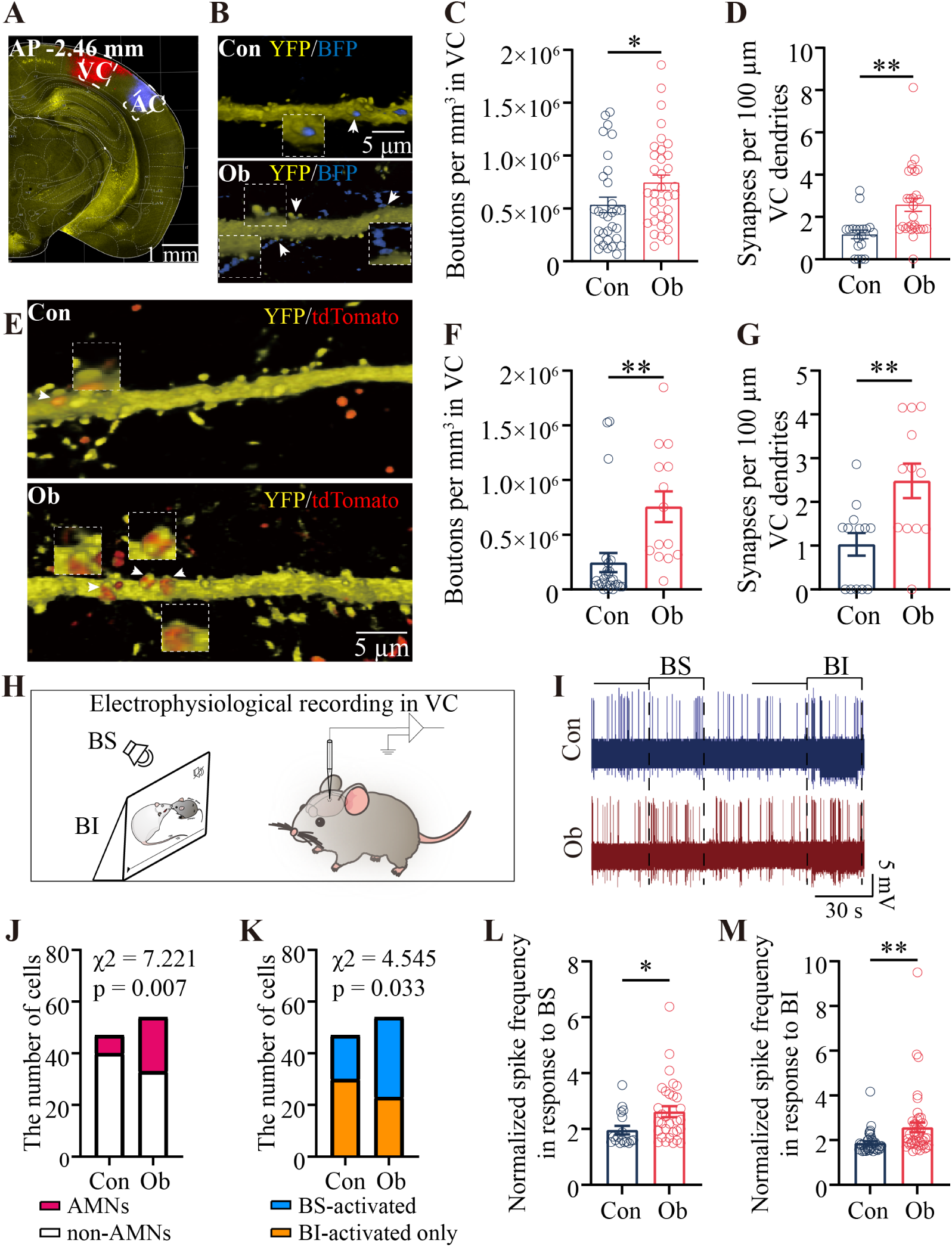
Psychological stress elevates the number of synaptic contacts and associative memory neurons in the VC. **A)** Confocal image illustrates the injection sites of AAV2/8-CMV-BFP in AC and AAV2/8-CMV-tdTomato in VC. **B)** BFP-labeled AC axonal boutons and their contacts on YFP-labeled dendritic spines of VC neurons from Con mouse (top panel) and Ob mouse (bottom panel). Images in white dash-line boxes show the amplification of synaptic contacts (1.68X) pointed by white arrows that denote synaptic contacts. **C)** The statistical analyses of axon boutons per mm^3^ in the VC projected from the AC in Con and Ob mice (one-way ANOVA). **D)** The statistical analyses of synaptic contacts per 100 m dendrite made between AC axon boutons and dendritic spines on VC neurons (one-way ANOVA). **E)** RFP-labeled VC axonal boutons and their contacts on YFP-labeled dendritic spines of VC neurons from Con mouse (top panel) and Ob mouse (bottom panel). Images in white dash-line boxes show the amplification of synaptic contacts (2.0X) pointed by white arrows that denote synaptic contacts. **F)** The statistical analyses of axon boutons per mm^3^ in the VC projected from the VC in Con and Ob mice (one-way ANOVA). **G)** The statistical analyses of synaptic contacts per 100 µm dendrite made between VC axon boutons and dendritic spines on VC neurons. **H)** A scheme shows *in vivo* electrophysiological recordings in VC as well as stimulations by battle image (BI) and battle sound (BS). **I)** The spikes at VC neurons spontaneously and evoked by BS and BI from Con mouse (blue) and Ob mouse (red). **J)** The statistical analyses about the numbers of associative memory neurons (AMNs, red in bar) that encode both BI and BS as well as of active neurons that encode either BI or BS (non-AMNs, white in bar) in the visual cortex from Con and Ob mice (chi-square test). **K)** The statistical analyses about the numbers of BS-activated neurons (blue in bar) and BI-activated neurons (orange in bar) in the visual cortex from Con and Ob mice (chi-square test). **L)** The statistical analyses about normalized spike frequencies in response to BS in VC neurons from Con and Ob mice (one-way ANOVA). **M)** The statistical analyses about normalized spike frequencies in response to BI in VC neurons from Con and Ob mice (one-way ANOVA). Abbreviations: AP: Anteroposterior; VC, visual cortex; AC, auditory cortex; BS: battle sound; BI: battle image. Data are presented as mean ± SEM. Two asterisks, p < 0.01; one asterisk, p < 0.05.

When the neurons in visual, auditory and medial frontal cortices receive two sources of synapse inputs, these cortical neurons are theoretically able to encode battle images and battle sounds in the resident/intruder paradigm, i.e., they are recruited to be associative memory neurons in morphology ^19,21,65^. Moreover, *in vivo* electrophysiological recordings in these cortices were conducted to examine whether these cortical neurons become able to encode battle sound and images. Spike frequencies were an index of neuronal activity strength. The evoked spikes on the background of spontaneous spikes were analyzed. Normalized spike frequency in response to one of stressful signals was calculated to be the ratio of the frequency of evoked spikes to the frequency of spontaneous spikes in 30 seconds before stimuli. If these ratios reached 1.5 or above, the recorded neurons were deemed as the responses to these signals. The neurons responded to one of two signals were called as active neurons. Auditory, visual and medial prefrontal cortical neurons in response to battle images and battle sound were presumably associative memory neurons in function, similar to previous studies^19,21,65,87,91–93^.

#### New synapses and associative memory neurons in the visual cortex

BFP-labelled axon boutons from auditory cortex to the visual cortex and their synapse contacts on visual cortical neurons appear more in observers with the fear memory (bottom panel in Figure 2B) than controls (top panel). Axon boutons per mm^3^ are 7.44x10^5^±0.73x10^5^ in observers (n=34 cubes from 5 mice) and 5.34x10^5^±0.74x 10^5^ in control group (n=31 cubes from 5 mice, p<0.05, ANOVA, Figure 2C). Synapse contacts per 100 μm dendrite on visual cortical neurons are 2.58±0.3 in observers (n=28 dendrites from 5 mice) and 1.17±0.2 in controls (n=19 dendrites from 5 mice, p=0.0015, ANOVA, Figure 2D). Furthermore, to the intramodal axon boutons and synapses in the visual cortex, the RFP-labelled axon boutons and their synapse contacts appear more in observers with fear memory (bottom panel in Figure 2E) than in controls (top panel). Axon boutons per mm^3^ are 7.57x10^5^± 1.41x10^5^ in observers (n=14 cubes from 4 mice) and 2.45x10^5^±0.87x10^5^ in controls (n=26 cubes from 4 mice, p=0.0024, ANOVA, Figure 2F). Synapse contacts per 100 μm dendrite on visual cortical neurons are 2.48±0.4 in observers (n=12 dendrites from 4 mice) and 1.03 ±0.26 in controls (n=13 dendrites from 4 mice, p=0.0048, ANOVA, Figure 2G). These data indicate that the synapse connections from the auditory cortex to the visual cortex and the synapse interconnections among visual cortical neurons are formed and increased in observational mice with the fear memory induced by psychological stress. The formation of new synapses is also supported by the upregulation of dendritic spines on visual cortical neurons from observational mice (Figure S5).

In the recording of unitary discharges at visual cortical neurons, battle sound and battle image in 30 seconds were given sequentially to observational mice with fear memory and control mice (Figure 2H). Some visual cortical neurons appear responding to both battle sound and image in observational mouse with fear memory (red trace in Figure 2I), but not control mouse (blue trace). The percentages of associative memory neurons in total active neurons were 38.89 % in observers (n=21/54 from 19 mice) and 14.89% in controls (n=7/47 from 15 mice; χ^2^=7.22, p=0.007, Chi-square test, Figure 2J). Such visual cortical neurons in observers do not respond to sounds in their living environments (Figure S2). This result indicates that the synapse innervations to visual cortical neurons from the auditory cortex recruit them to encode battle sounds and battle images, or associative memory neurons, for the joint storage of fear signals. This indication is granted by a result that the number of visual cortical neurons in response to battle sounds is more in observers than in controls (χ^2^=4.55, p=0.033, Figure 2K). In addition, normalized spike frequencies at active visual cortical neurons in response to battle sounds are 2.62 ±0.20 in observers (n=31 cells from 19 mice) and 1.96±0.15 in control (n=17 cells from 15 mice, p=0.026, ANOVA, Figure 2L), indicating that the synapse innervations from the auditory cortex (Figure 2B-D) enhance the excitatory activity of visual cortical neurons. Normalized spike frequencies at active visual cortical neurons in response to battle images are 2.57±0.22 in observers (n=44 cells from 19 mice) and 1.87± 0.08 in controls (n=37 cells from 15 mice, p=0.006, ANOVA, Figure 2M), indicating that the synapse innervations raised within intramodal visual cortex (Figure 2E-G) enhance the excitatory activity of visual cortical neurons.

#### New synapses and associative memory neurons in auditory cortex

RFP-labelled axon boutons from the visual cortex to the auditory cortex and their synapse contacts appear more in observational mice with fear memory (bottom panel in Figure 3A) than controls (top panel). Axon boutons per mm^3^ are 4.76x10^5^± 0.66x10^5^ in observer (n=35 cubes from 5 mice) and 2.54x10^5^±0.27 x10^5^ in control (n=25 cubes from 5 mice, p=0.008, ANOVA, Figure 3B). Synapse contacts per 100 μm dendrite on auditory cortical neurons are 1.68±0.18 in observers (n=35 dendrites from 5 mice) and 0.83±0.18 in controls (n=16 dendrites from 5 mice, p=0.0049, ANOVA, Figure 3C). Furthermore, to intramodal axon boutons and synapses in the auditory cortex, BFP-labelled synapse contacts appear more in observers with fear memory (bottom panel in Figure 3D) than in controls (top panel). Axon boutons per mm^3^ are 3.77x10^6^±0.59x10^6^ in observers (n=26 cubes from 5 mice) and 3.26x10^6^±0.35x10^6^ in controls (n=43 cubes from 6 mice, p=0.43, ANOVA, Figure 3E). Synapse contacts per 100 μm dendrite on auditory cortical neurons are 2.61±0.33 in observers (n=24 dendrites from 9 mice) and 1.46±0.23 in controls (n=31 dendrites from 5 mice, p=0.0049, ANOVA, Figure 3F). Thus, the synapse connections from the visual cortex to the auditory cortex as well as the synapse interconnections among auditory cortical neurons are formed and increased in observational mice with stress-induced fears. The formation of new synapses is also supported by the change of dendritic spines on auditory cortical neurons from observational mice (Figure S5).

**Figure 3.**
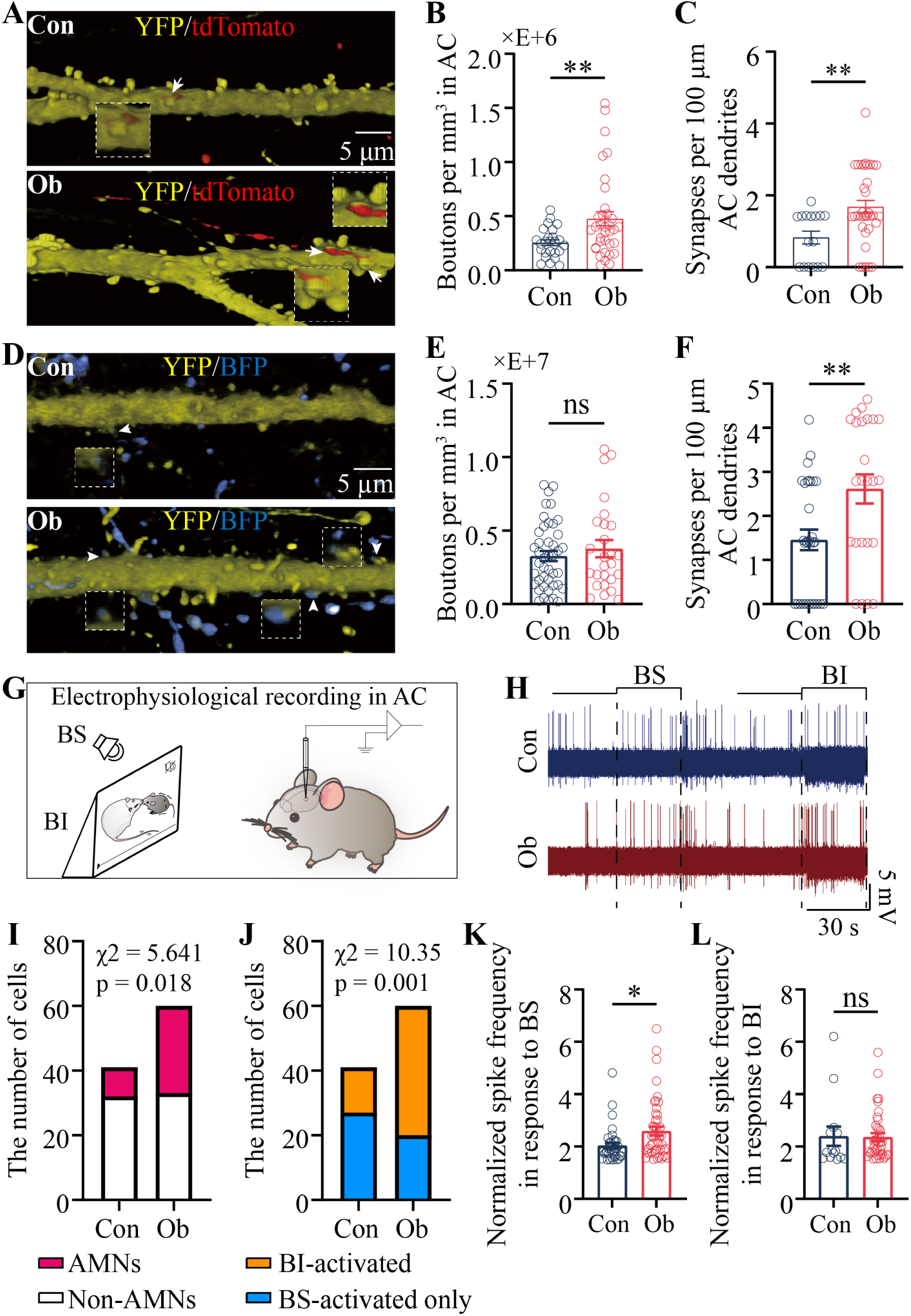
Psychological stress elevates the number of synaptic contacts and associative memory neurons in the AC. **A)** RFP-labeled VC axon boutons and their contacts on YFP-labeled dendritic spines of AC neurons from Con mouse (top panel) and Ob mouse (bottom panel). Images in white dash-line boxes show the amplification of synaptic contacts (1.68X) pointed by white arrows that denote synaptic contacts. **B)** The statistical analyses of axon boutons per mm^3^ in the AC projected from the VC in Con and Ob mice (one-way ANOVA). **C)** The statistical analyses of synaptic contacts per 100 µm dendrite made between VC axon boutons and dendritic spines on AC neurons (one-way ANOVA). **D)** BFP-labeled AC axon boutons and their contacts on YFP-labeled dendritic spines of AC neurons from Con mouse (top panel) and Ob mouse (bottom panel). Images in white dash-line boxes show the amplification of synaptic contacts (2.0X) pointed by white arrows that denote synaptic contacts. **E)** The statistical analyses of axon boutons per mm^3^ in the AC projected from the AC in Con and Ob mice (one-way ANOVA). **F)** The statistical analyses of synaptic contacts per 100 µm dendrite made between AC axon boutons and dendritic spines on AC neurons. **G)** A scheme shows *in vivo* electrophysiological recordings in AC as well as stimulations by battle image (BI) and battle sound (BS). **H)** The spikes at AC neurons spontaneously and evoked by BS and BI from Con mouse (blue) and Ob mouse (red). **I)** The statistical analyses about the numbers of associative memory neurons (AMNs, red in bar) that encode both BI and BS as well as of active neurons that encode either BI or BS (non-AMNs, white in bar) in the auditory cortex from Con and Ob mice (chi-square test). **J)** The statistical analyses about the numbers of BI-activated neurons (orange in bar) and BS-activated neurons (blue in bar) in the auditory cortex from Con and Ob mice (chi-square test). **K)** The statistical analyses about normalized spike frequencies in response to BS in AC neurons from Con and Ob mice (one-way ANOVA). **L)** The statistical analyses about normalized spike frequencies in response to BI in AC neurons from Con and Ob mice (one-way ANOVA). Abbreviations: AP: Anteroposterior; VC, visual cortex; AC, auditory cortex; BS: battle sound; BI: battle image. Data are presented as mean ± SEM. Two asterisks, p < 0.01; one asterisk, p < 0.05; ns, p >0.05.

The *in vivo* electrophysiological recordings in the auditory cortex were used to examine whether auditory cortical neurons become able to encode battle sound and battle image (Figure 3G). Auditory cortical neurons respond to both battle sounds and images in observational mouse with fear memory (red trace in Figure 3H), but not in control mouse (blue trace). The percentages of associative memory neurons in total active neurons were 45% in observers (n=27/60 from 19 mice) and 22% in controls (n=9/41 from 14 mice; χ^2^= 5.641, p=0.018, Chi-square test, Figure 3I). These auditory cortical neurons in observers do not respond to flower images (Figure S2). This result indicates that the synapse inputs to auditory cortical neurons from the visual cortex recruit them as associative memory neurons that encode battle sounds and images for the joint storage of fear signals. This indication is supported by the result that the number of auditory cortical neurons in response to battle images appears more in observers than in controls (χ^2^=10.35, p=0.001, Figure 3J). Furthermore, normalized spike frequencies at active auditory cortical neurons in response to battle sounds are 2.59±0.17 in observers (n=47 cells from 15 mice) and 2.04±0.11 in controls (n=36 cells from 14 mice, p=0.01, ANOVA, Figure 3K), i.e., the synapse innervations raised in the intramodal auditory cortex (Figure 3D) strengthen the excitatory activity of auditory cortical neurons. However, normalized spike frequencies at active auditory cortical neurons in response to battle images are 2.36±0.15 in observers (n=40 cells in 15 mice) and 2.4±0.37 in controls (blue bar, n=14 cells from 15 mice, p=0.92, ANOVA, Figure 3K), implying that the synapse innervations increased from the cross-modal visual cortex (Figure 3A-C) are not sufficient to enhance the excitatory activity of auditory cortical neurons.

#### New synapses and associative memory neurons in the medial prefrontal cortex

As the medial prefrontal cortex locates at the downstream of sensory cortices in neuronal signal flows ^94,95^ and prefrontal cortical neurons can be recruited as associative memory cells to encode physiological signals ^64,93,96,97^, these medial prefrontal cortical neurons may be recruited as associative memory neurons to encode the fear signals in the psychological stress based on receiving convergent synapse innervations from visual and auditory cortices.

To study axon boutons and their synapse contacts in the medial prefrontal cortex crossmodally from visual and auditory cortices, we microinjected 0.3 μl AAV-CMV-tdTomato in the visual cortex and 0.3 μl AAV-CMV-BFP in the auditory cortex (top panel in Figure 4A). RFP-labelled axon boutons, BFP-labelled axon boutons and synapse contacts made by these axon inputs appear more in the medial prefrontal cortex in observational mice with fear memory (bottom panel in Figure 4B) than in controls (top and middle panels). Axon boutons per mm^3^ in the medial prefrontal cortex are 1.82x10^6^±0.4x10^6^ in observers (n=27 cubes from 5 mice) and 0.63x10^6^±0.14x10^6^ in controls (n=36 cubes from 5 mice, p=0.003, ANOVA, Figure 4C). Synapse contacts per 100 μm dendrite onto medial prefrontal cortical neurons are 2.18±0.2 in observers (n=31 dendrites from 5 mice) and 0.81±0.16 in controls (n=22 dendrites from 5 mice, p<0.0001, ANOVA, Figure 4D). Moreover, to examine intramodal axon boutons and their synapse contacts in the medial prefrontal cortex, we microinjected 0.3 μl AAV-CMV-EGFP in this area (bottom panel in Figure 4A). GFP-labelled intramodal axon boutons and the synapse contacts appear more in observers with fear memory (bottom panel in Figure 4E) than in controls (top panel). Axon boutons per mm^3^ are 1.07x10^7^±0.39x10^7^ in observers (n=12 cubes from 5 mice) and 0.27x10^7^± 0.1x10^7^ in controls (n=16 cubes from 5 mice, p=0.035, ANOVA, Figure 4F). Synapse contacts per 100 μm dendrite on medial prefrontal cortical neurons are 4.42±0.43 in observers (n=10 dendrites from 4 mice) and 2.99±0.45 in controls (n=9 dendrites from 4 mice, p=0.036, ANOVA, Figure 4G). Therefore, the convergent synapse innervations on medial prefrontal cortical neurons from visual and auditory cortices and the synapse interconnections among medial prefrontal cortical neurons are formed and increased in observational mice with fears.

**Figure 4.**
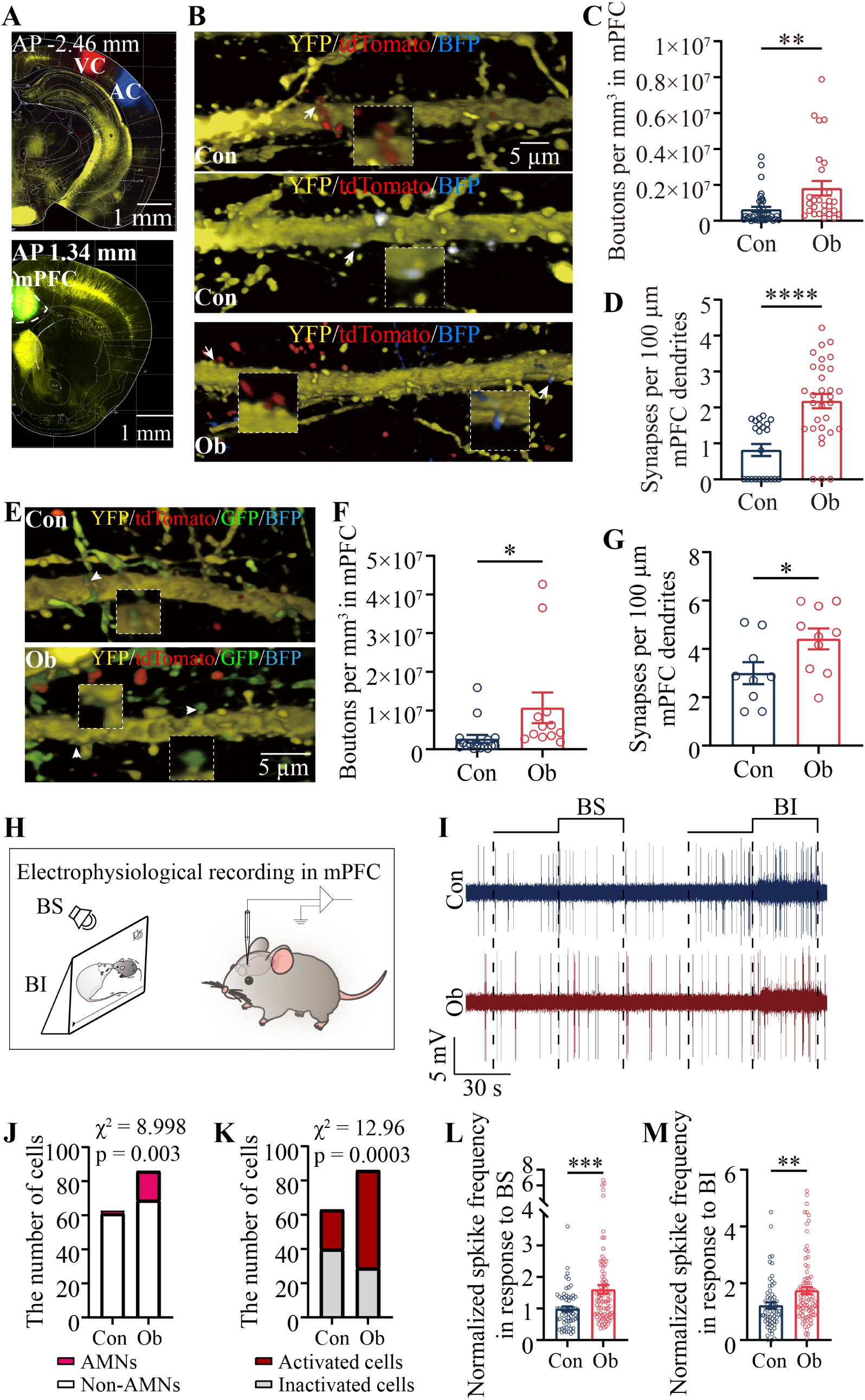
Psychological stress elevates the number of synaptic contacts and associative memory neurons in the mPFC. **A)** Confocal image demonstrates the injection sites of AAV2/8-CMV-BFP in the AC, AAV2/8-CMV-tdTomato in the VC (top panel) and AAV2/8-CMV-GFP in mPFC (bottom panel). **B)** RFP-labeled VC axon boutons, BFP-labeled AC boutons and their contacts on YFP-labeled dendritic spines of mPFC neurons from Con mouse (top panel and middle panel) and Ob mouse (bottom panel). Images in white dash-line boxes show the amplification of synaptic contacts (2.0X) pointed by white arrows that denote synaptic contacts. **C)** The statistical analyses of axon boutons per mm^3^ in the mPFC projected from the AC and VC in Con and Ob mice (one-way ANOVA). **D)** The statistical analyses of synaptic contacts per 100 µm dendrite made among AC/VC axon boutons and dendritic spines on mPFC neurons (one-way ANOVA). **E)** GFP-labeled mPFC axon boutons and their contacts on YFP-labeled dendritic spines of mPFC neurons from Con mouse (top panel) and Ob mouse (bottom panel). Images in white dash-line boxes show the amplification of synaptic contacts (2.0X) pointed by white arrows that denote synaptic contacts. **F)** The statistical analyses of axon boutons per mm^3^ in the mPFC projected from the mPFC in Con and Ob mice (one-way ANOVA). **G)** The statistical analyses of synaptic contacts per 100 µm dendrite made between mPFC axon boutons and dendritic spines on mPFC neurons. **H)** A scheme shows *in vivo* electrophysiological recordings in mPFC as well as stimulations by battle image (BI) and battle sound (BS). **I)** The spikes at mPFC neurons spontaneously and evoked by BS and BI from Con mouse (blue) and Ob mouse (red). **J)** The statistical analyses about the numbers of associative memory neurons (AMNs, red in bar) that encode both BI and BS as well as of non-associative memory neurons (non-AMNs, white in bar) that encode BI, BS or neither signals in the mPFC from Con and Ob mice (chi-square test). **K)** The statistical analyses about the numbers of activated neurons (red in bar) and inactivated neurons that encode neither BI nor BS (grey in bar) in the mPFC from Con and Ob mice (chi-square test). **L)** The statistical analyses about normalized spike frequencies in response to BS in mPFC neurons from Con and Ob mice (one-way ANOVA). **M)** The statistical analyses about normalized spike frequencies in response to BI in mPFC neurons from Con and Ob mice (one-way ANOVA). Abbreviations: AP: Anteroposterior; mPFC: medial prefrontal cortex; VC, visual cortex; AC, auditory cortex; BS: battle sound; BI: battle image. Data are presented as mean ± SEM. Four asterisks, p < 0.0001; three asterisks, p < 0.001; two asterisks, p < 0.01; one asterisk, p < 0.05.

The *in vivo* electrophysiological recordings in the medial prefrontal cortex were used to examine whether medial prefrontal cortical neurons become able to encode stressful signals (Figure 4H). Some medial prefrontal cortical neurons respond to battle sounds and images in observational mouse with fear memory (red trace in Figure 4I), but not control one (blue trace). The percentages of associative memory neurons in the total neurons were 19.77 % in observers (n=17/86 from 10 mice) and 3.17% in controls (n=2/63 from 7 mice; χ^2^= 9.0, p=0.003, Chi-square test, Figure 4J). Medial prefrontal cortical neurons in observers do not respond to the environmental sound and flower image (Figure S2). These data indicate that synapse innervations on medial prefrontal cortical neurons from visual and auditory cortices recruit them to encode battle sound and battle image, or associative memory neuron, for the acquired fear induced by psychological stress. This indication is granted by a result that the number of medial prefrontal cortical neurons in response to battle images or battle sounds is more in observers than controls (χ^2^=12.96, p=0.0003, Figure 4K). Furthermore, normalized spike frequencies at medial prefrontal cortical neurons in response to battle sound are 1.61±0.14 in observers (n=86 cells from 10 mice) and 0.99±0.08 in controls (n=63 cells from 7 mice, p=0.0005, ANOVA, Figure 4L). Normalized spike frequencies at medial prefrontal cortical neurons in response to battle images are 1.74±0.13 in observers (n=86 cells from 10 mice) and 1.22±0.11 in controls (n=63 cells from 7 mice, p=0.0035, ANOVA, Figure 4M). These data indicate that the synapse innervations on medial prefrontal cortical neurons increased from cross-modal visual and auditory cortices (Figure 4B-D) and among intramodal neurons (Figure 4E-G) strengthen the excitatory activity of medial prefrontal cortical neurons.

Both morphological and electrophysiological data validate that psychological stress recruits the associative memory neurons in visual, auditory and medial prefrontal cortices. The formation of new synapse innervations in these cortices crossmodally from their interconnected areas in the mice with fear memory recruits cortical neurons to be associative memory neurons that encode fear signals. The synapse innervations from cross-modal cortical neurons as well as the synapse interconnection among intramodal neurons strengthen the spike-encoding of associative memory neurons and active cortical neurons.

## Kinesin family member-1A is required for recruiting associative memory neurons to encode fears

In terms of the mechanisms underlying the interconnections among visual, auditory and medial prefrontal cortices and the recruitment of associative memory neurons to encode the phobia, the molecules related to the axon growth and transportation are presumably involved. Kinesins are motor proteins for the intracellular transportation of vesicles in axons and receptors in dendrites as well as the function of synapses ^98–109^, which are relevant to learning and memory ^110,111^. Kinesins are also highly expressed in the medial prefrontal cortex of observational mice (Figure S3). We hypothesized that kinesins might be required for the elongation of axons, the enlargement of dendritic spines and the formation of new synapses to recruit fear memory neurons. The knockdown of kinesin family member-1a (KiF1a) in the medial prefrontal cortex was conducted to examine the requirement of KiF1a for these processes. A rationale for using *KiF1a* knockdown to suppress the recruitment of associative memory neurons is based on the axon projection for their recruitment ^19,21,65,87,89,91^.

AAV-carried RNAi specifically to *KiF1a* (AAV-U6-shKiF1a-GFP) was injected in medial prefrontal cortices of observational mice five days before their facing to a resident/intruder paradigm (bottom panel in Figure 5A). This shRNA has been proven to reduce KiF1a expression in the medial prefrontal cortex (Figure S3). shRNA scramble control was injected in another group of observational mice. The experiments in *KiF1a* knockdown (*KiF1a*-KD) and scramble controls were conducted by behavior tasks, neural tracing and electrophysiology *in vivo* applied in Figures 1-4. The essential roles of KiF1a in the onset of fear memory, the formation of new synapse innervations and the recruitment of associative memory neurons would be ensured if these processes were lowered in observational mice with *KiF1a* knockdown.

**Figure 5.**
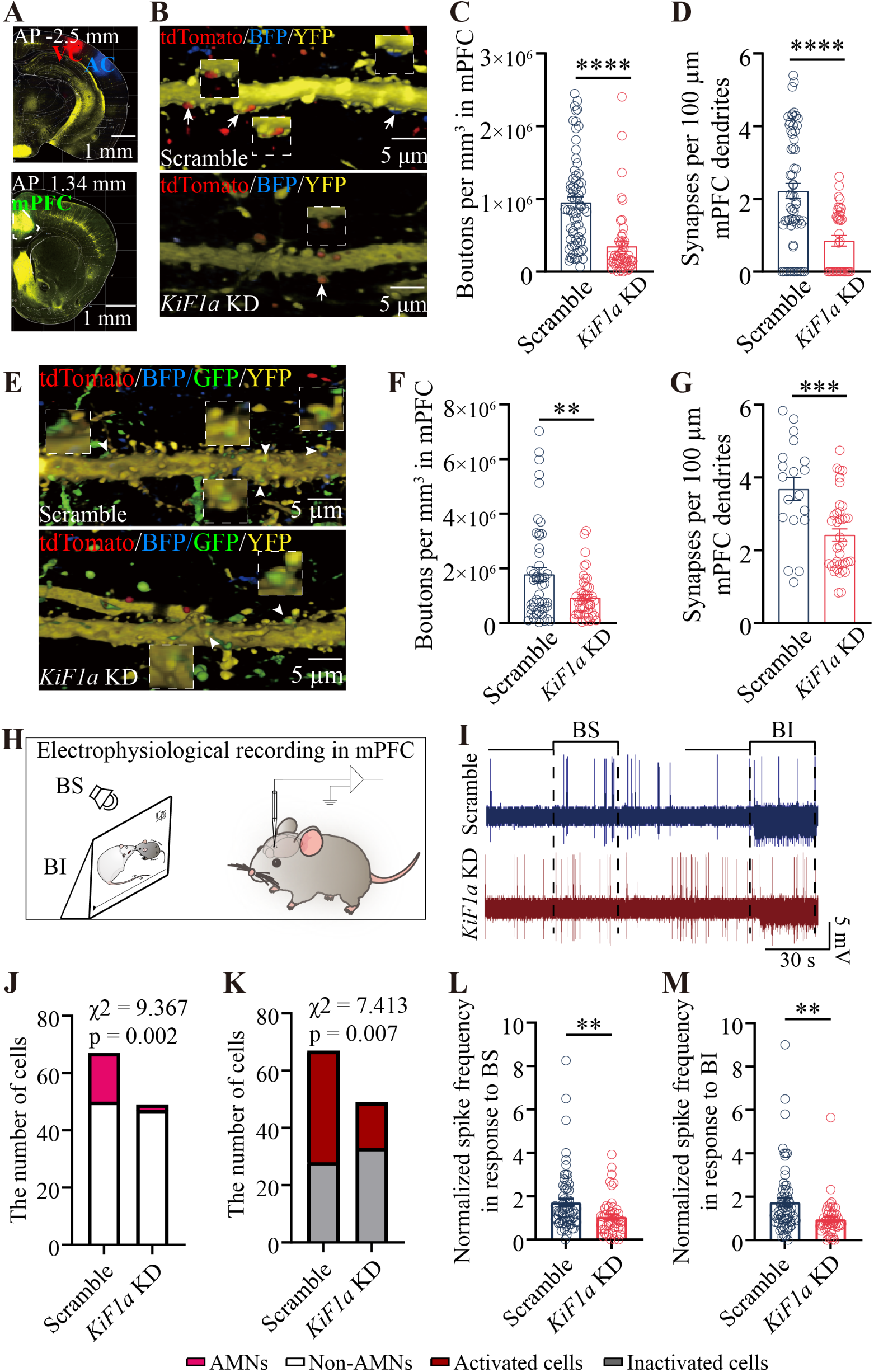
*KiF1a* RNAi in mPFC precludes the stress-induced recruitments of axon projection, synapse formation and associative memory neurons. **A)** Confocal image illustrates the injection sites of AAV2/8-CMV-BFP in AC, AAV2/8-CMV-tdTomato in VC (top panel) and AAV2/9-U6-shRNA-GFP in mPFC (bottom panel). **B)** RFP-labeled VC axon boutons, BFP-labeled AC boutons and their contacts on YFP-labeled dendritic spines of mPFC neurons from scramble mouse (top panel) and *KiF1a*-KD mouse (bottom panel). Images in white dash-line boxes show the amplification of synaptic contacts (1.2X) pointed by white arrows that denote synaptic contacts. **C)** The statistical analyses of axon boutons per mm^3^ in the mPFC projected from the AC and VC in scramble and *KiF1a*-KD mice (one-way ANOVA). **D)** The statistical analyses of synaptic contacts per 100 µm dendrite made among AC, VC axon boutons and dendritic spines on mPFC neurons (one-way ANOVA). **E)** GFP-labeled mPFC axon boutons and their contacts on YFP-labeled dendritic spines of mPFC neurons from scramble mouse (top panel) and *KiF1a*-KD mouse (bottom panel). Images in white dash-line boxes show the amplification of synaptic contacts (2.3X) pointed by white arrows that denote synaptic contacts. **F)** The statistical analyses of axon boutons per mm^3^ in the mPFC projected from the mPFC in scramble and *KiF1a*-KD mice (one-way ANOVA). **G)** The statistical analyses of synaptic contacts per 100 µm dendrite made between mPFC axon boutons and dendritic spines on mPFC neurons. **H)** A scheme shows *in vivo* electrophysiological recordings in mPFC as well as stimulations by battle image (BI) and battle sound (BS). **I)** The spikes at mPFC neurons spontaneously and evoked by BS and BI from scramble mouse (blue) and *KiF1a*-KD mouse (red). **J)** The statistical analyses about the numbers of associative memory neurons (AMNs, red in bar) that encode both BI and BS as well as of non-associative memory neurons (non-AMNs, white in bar) that encode BI, BS or neither signals in the mPFC cortex from scramble and *KiF1a*-KD mice (chi-square test). **K)** The statistical analyses about the numbers of active neurons (red in bar) and inactive neurons that encode neither BI nor BS (grey in bar) in the mPFC cortex from scramble and *KiF1a*-KD mice (chi-square test). **L)** The statistical analyses about normalized spike frequencies in response to BS in mPFC neurons from scramble and *KiF1a*-KD mice (one-way ANOVA). **M)** The statistical analyses about normalized spike frequencies in response to BI in mPFC neurons from scramble and *KiF1a*-KD mice (one-way ANOVA). Abbreviations: AP: Anteroposterior; mPFC: medial prefrontal cortex; VC, visual cortex; AC, auditory cortex; BS: battle sound; BI: battle image. Data are presented as mean ± SEM. our asterisks, p < 0.0001; three asterisks, p < 0.001; two asterisks, p < 0.01.

AAV-CMV-tdTomato and AAV-CMV-BFP were microinjected in the visual cortex and the auditory cortex, respectively, when AAV-U6-shKiF1a-GFP was injected in the medial prefrontal cortex (Figure 5A). RFP-labelled axon boutons from the visual cortex, BFP-labelled axon boutons from the auditory cortex and such axon-made synapse contacts on dendritic spines of medial prefrontal cortical neurons (yellow) appear lower in observational mice with *KiF1a*-KD (bottom panel in Figure 5B) than scramble control mice (top panel). Axon boutons per mm^3^ in the medial prefrontal cortex are 3.48x10^5^±0.65x 10^5^ in *KiF1a*-KD group (n=51 cubes from 5 mice) and 9.52x10^5^± 0.76x10^5^ in scramble controls (n=69 cubes from 5 mice, p<0.0001, ANOVA, Figure 5C). Synapse contacts per 100 μm dendrite on medial prefrontal cortical neurons are 0.85±0.15 in *KiF1a*-KD (n=36 dendrites from 5 mice) and 2.22±0.21 in scrambles (n=62 dendrites from 5 mice, p<0.0001, ANOVA, Figure 5D). In addition, intramodal axon boutons and their synapse contacts in the medial prefrontal cortex were examined by GFP-mediated tracing (AAV-U6-shKiF1a-GFP, bottom panel in Figure 5A). GFP-labelled intramodal axon boutons and their synapse contacts appear lower in observers with *KiF1a*-KD (bottom panel in Figure 5E) than in scrambles (top panel). Axon boutons per mm^3^ are 9.25x10^5^±1.16x10^5^ in *KiF1a*-KD (n=46 cubes of 5 mice) and 17.74x10^5^±2.49x10^5^ in scrambles (n=50 cubes from 9 mice, p<0.01, ANOVA, Figure 5F). Synapse contacts per 100 μm dendrite on medial prefrontal cortical neurons are 2.42±0.17 in *KiF1a*-KD (n=38 dendrites from 5 mice) and 3.68±0.32 in scrambles (n=19 dendrites from 5 mice, p=0.0003, ANOVA, Figure 5G). KiF1a is required for the formation of new cross-modal and intramodal synapse innervations and for the recruitment of associative memory neurons in the medial prefrontal cortex induced by psychological stress.

The *in vivo* electrophysiological recordings in the medial prefrontal cortex were done to examine the functional effectiveness of KiF1a on the recruitment of associative memory neurons in the medial prefrontal cortex (Figure 5H). Medial prefrontal cortical neurons respond to battle sound and images in scramble control mouse (blue trace in Figure 5I), but not observer with *KiF1a*-KD (red trace). The percentages of associative memory neurons in all recorded neurons were 4.08% in *KiF1a*-KD (n=2/49 from 7 mice) and 25.37% in scramble control (n=17/67 from 7 mice; χ^2^= 9.37, p=0.002, Chi-square test, Figure 5J). The downregulation of synapse innervations on medial prefrontal cortical neurons from the visual and auditory cortices precludes them to encode battle sound and images in the phobia induced by psychological stress. This indication is supported by the data that the number of medial prefrontal cortical neurons in response to battle images or battle sounds is more in scrambles than in *KiF1a*-KD (χ^2^=7.41, p=0.007, Figure 5K). In addition, normalized spike frequencies at medial prefrontal cortical neurons in response to battle sound are 1.05±0.13 in *KiF1a*-KD (n=49 cells from 7 mice) and 1.71±0.17 in scrambles (n=67 cells from 7 mice, p=0.0047 ANOVA, Figure 5L). Normalized spike frequencies at medial prefrontal cortical neurons in response to battle images are 0.96±0.12 in *KiF1a*-KD (n=49 cells from 7 mice) and 1.75±0.19 in scrambles (n=67 cells from 7 mice, p=0.002, ANOVA, Figure 5M). Thus, the downregulation of synapse innervations from cross-modal and intramodal cortices onto medial prefrontal cortical neurons that encode battle sound and battle images weakens the excitatory activity of medial prefrontal cortical neurons.

In the experiment of behavior tasks, the observational mice with KiF1a-KD appear not to avoid the resident mouse (right panel in Figure 6A), compared with scramble control mice (left panel). The interaction time (seconds) in response to a resident mouse is 125.6±8.3 in *KiF1a*-KD mice (n=16) and 88.96±8.53 in scrambles (n=14; p=0.0047, ANOVA, Figure 6B). Furthermore, the interaction time in response to battle sound appears lower in scramble controls (left panel in Figure 6C) than *KiF1a*-KD mice (right panel). The interaction time (second) in response to battle sound is 121.1±8.62 in *KiF1a*-KD mice (n=17) and 88.26±5.52 in scramble mice (n=16, p<0.01, ANOVA, Figure 6D). It is noteworthy that the mice with injections of AAV-U6-sh*KiF1a*-GFP and scramble controls in medial prefrontal cortices do not show fear to CD1 resident mouse, battle sound, housing environment sound and mouse surrogate before the experiments as well as not fear to housing environment sound and mouse surrogates after the psychological stress (Figure S4). Therefore, *KiF1a* knockdown in the medial prefrontal cortices of observational mice precludes the retrieval of fear memories to resident mice and battle sounds.

**Figure 6.**
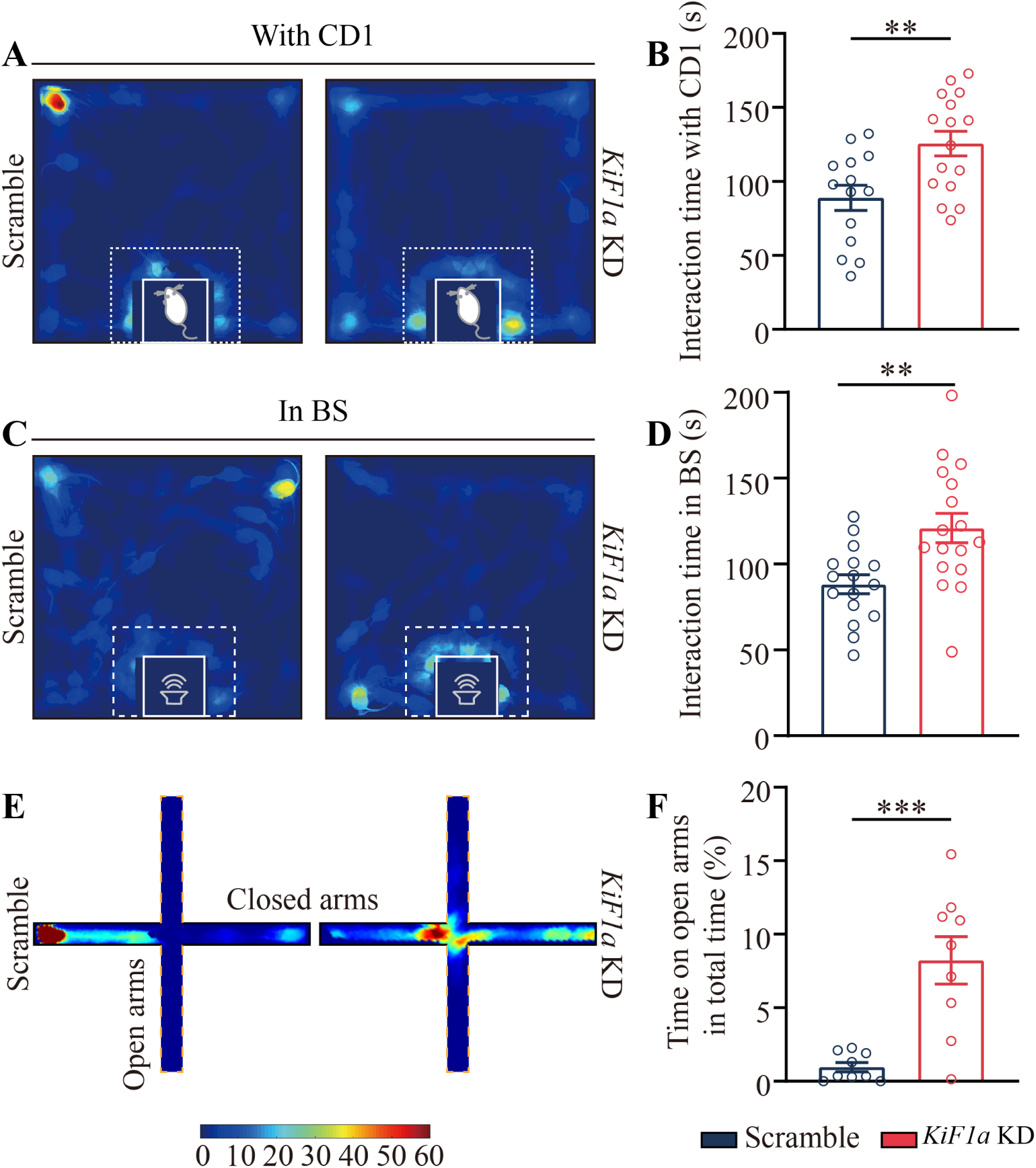
*KiF1a* RNAi in mPFC attenuates stress-induced fear memory and anxiety-like behavior. **A)** Heat maps illustrate the social interaction test for C57 scramble and *KiF1a-*KD mice in face to a CD1 resident mouse. **B)** A bar graph with samples’ dots shows the interaction time of Ob plus *KiF1a-*KD and scramble mice with CD1 resident. **C)** Heat maps show the social interaction test for Ob plus *KiF1a-*KD and scramble mice in response to battle sound. **D)** A bar graph with samples’ dots shows the interaction time of Ob plus *KiF1a-*KD and scramble mice with battle sound. **E)** Heat maps illustrate the elevated-plus maze test for Ob plus scramble and *KiF1a* KD mice. **F)** A bar graph with samples’ dots for show the time spent on open arms in the total time on the elevated-plus maze. Abbreviations: KD: knock down; BS, battle sound. Data are presented as mean ± SEM and analyzed by one-Way ANOVA. Three asterisks, p<0.001; two asterisks, p<0.01

The influence of *KiF1a* knockdown on anxiety-like behaviors of observational mice was examined by using the EPM. The stay time on the plate of open arms in the EPM appears longer in observational mice with *KiF1a*-KD (right panel in Figure 6E) than scramble control mice (left panel). The ratios of the stay time on the plate of the open arms to the total time in the EPM are 8.22±1.61% in *KiF1a*-KD mice (n=9) and 0.96±0.31% in scramble control mice (n=9, p=0.0004, ANOVA, Figure 6F). *KiF1a* knockdown in medial prefrontal cortices of observational mice precludes their anxiety-like behaviors induced by psychological stress. These data indicate that KiF1a in the medial prefrontal cortex plays an essential role in stress-induced fear memory and anxiety.

## Associative memory neurons in the medial prefrontal cortex are secondary in nature

Psychological stress recruits associative memory neurons that encode stressful signals in visual, auditory and medial prefrontal cortices. As the medial prefrontal cortex anatomically and functionally locates at the downstream of visual and auditory cortices, the associative memory neurons in sensory cortices might be primary, and the associative memory neurons in the medial prefrontal cortex might be secondary ^64,96,97^. The assumption can be proven if the encoding of stress signals at associative memory neurons in the medial prefrontal cortex is prevented by downregulating the recruitment of associative memory neurons in sensory cortices as well as blocked by inhibiting the activity of sensory cortical neurons. The prevention of recruiting the associative memory neurons in visual and auditory cortices was conducted by the *KiF1a* knockdown in these areas. The activity blockage of visual and auditory cortical neurons was conducted by CNQX and D-AP5, the antagonists of inotropic glutamate receptors, in these areas.

The KiF1a knockdown in visual and auditory cortices of observational mice (Figure 7A) appears to prevent the responses of medial prefrontal cortical neurons to battle sounds and images (red trace in Figure 7B), in comparison with scramble controls (blue trace). The percentages of associative memory neurons in total recorded neurons were 7.46% in KiF1a-KD group (n=5/67 from 4 mice) and 24.1% in scramble group (n=13/54 from 4 mice; χ^2^= 6.52, p= 0.01, Chi-square test, Figure 7C). The percentages of active neurons in total neurons were 40.3% in KiF1a-KD group (n=27/67 from 4 mice) and 62.96% in scramble controls (n= 34/54 from 4 mice; χ^2^= 6.14, p=0.01, Chi-square test, Figure 7D). Moreover, the blockages of ionotropic glutamate receptors in the visual and auditory cortices of observational mice by 10 μM CNQX and 40 μM D-AP5 (Figure 7E) appear to attenuate the responses of medial prefrontal cortical neurons to battle sounds and images (Figure 7F). The ratios of normalized spike frequencies in responses to battle sounds (Figure 7G) and battle images (Figure 7H) are attenuated to less 1.5. These suppressed responses of associative memory neurons to stress signals in the medial prefrontal cortex by downregulating the activities of visual and auditory cortices indicate that the associative memory neurons in sensory cortices are primary and the associative memory neurons in the medial prefrontal cortex are secondary, which are encode fear and anxiety to be called as anxiety-coding neurons.

**Figure 7.**
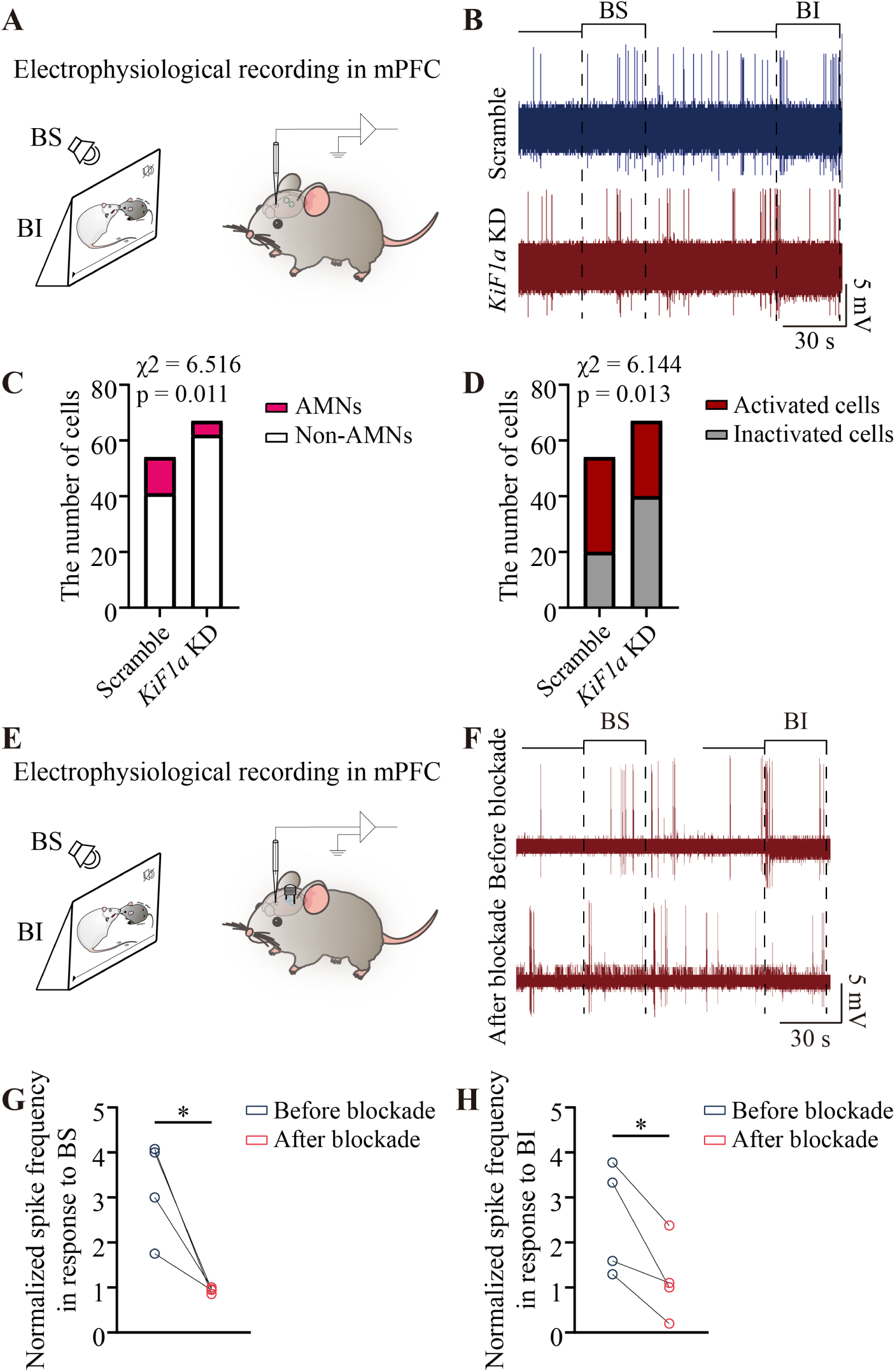
The inhibition of VC and AC neurons impairs the activity of associative memory neurons in mPFC. **A)** A scheme shows *in vivo* electrophysiological recordings in the mPFC and the stimulations of battle sound (BS) and battle image (BI). Green dashed circles on the tope of mouse head show AAV-shRNA (scramble) or AAV-sh*KiF1a* are simultaneously injected into VC and AC. **B)** The spikes at mPFC associative memory neurons spontaneously and evoked by BS and BI from scramble mouse (blue) and *KiF1a*-KD mouse (red). **C)** The statistical analyses about the numbers of associative memory neurons (AMNs, red in bar) that encode both BI and BS as well as of non-associative memory neurons (non-AMNs, white in bar) that encode BI, BS or neither signals in the mPFC from scramble and *KiF1a*-KD mice (chi-square test). **D)** The statistical analyses about the numbers of active neurons (red in bar) and inactive neurons that encode neither BI nor BS (grey in bar) in the mPFC from scramble and *KiF1a*-KD mice (chi-square test). **E)** A diagram shows *in vivo* electrophysiological recordings in the mPFC and stimulations of battle sound (BS) and battle image (BI). Blue dashed circles on the top of mouse head show VC and AC blockade by CNQX and DAP5. **F)** The spikes at mPFC associative memory neurons spontaneously and evoked by BS and BI before and after AC and VC blockade. **G)** The statistical analyses about normalized spike frequencies in response to BS in mPFC associative memory neurons before and after AC and VC blockade (paired *t*-test). **H)** The statistical analyses about normalized spike frequencies in response to BI in mPFC associative memory neurons before and after AC and blockade (paired *t*-test). Abbreviation: KD: knock down; BS: battle sound; BI: battle image. One asterisk, p < 0.05.

## Discussion

The psychological stress by watching a resident/intruder paradigm induces the fear memory specific to stressful scenes and anxiety-like behavior (Figure 1). In the mice with this acquired fear, the synapse interconnections between visual and auditory cortices and their convergent synapses on medial prefrontal cortical neurons are newly formed and functionally active. Some neurons in these cortical areas become able to encode stress signals including battle sound and battle image (Figures 2-4). Thus, psychological stress recruits associative memory neurons in neural pathways from visual and auditory cortices to the medial prefrontal cortex convergently (Figures 2-4) that encode psychosis. The recruitment of associative memory neurons and the onset of acquired fear are precluded by KiF1a knockdown, one of motor proteins (Figures 5-6). Therefore, psychological stress recruits visual, auditory and medial prefrontal cortical neurons as the associative memory neurons that encode fear-relevant psychosis by the KiF1a-mediated transportation of intracellular molecules and organelles.

In terms of the mechanisms for the recruitment of associative memory neurons in auditory, visual and medial prefrontal cortices to encode acquired fear, our interpretations are given below. The stress signals in the battle scenes of resident/intruder paradigm were dissected and sensed by visual and auditory systems, which activate visual and auditory cortical neurons. Based on the rule of coactivity together and interconnection together ^19,91,92^, the simultaneously intensive activities of these cortical neurons instigate biological cascades from higher frequency action potentials to epigenetic events ^19,87–90^, which then trigger axon elongation ^19,87,89^ and enhance KiF1a-mediated intracellular transportation ^98–104,106^. These neural events facilitate the projection of the axons to their active target areas as well as the transportation of presynaptic vesicles and building molecules for axon innervations to their target neurons ^98,100,103^(Figures 2-4). These biological cascades in the coactive postsynaptic neurons may induce the expansion of dendritic arbors, the transportation of intra-dendritic molecules ^106–109,112,113^ and the increase of dendritic spines ^21^ (Figure S5). The upregulations of presynaptic axon arbors and postsynaptic dendritic spines may work coordinately for the formation of new synapses. When presynaptic boutons and postsynaptic spines contact, these new synapses need be anchored by synapse-linkage proteins, neuroligins and neurexin ^21,65,114–117^. Whether the expansion of dendrites and spines needs KiF1a-mediated intracellular transportation about postsynaptic receptors and synapse-building molecules remains to be examined. New axon projection, spine increase and synapse formation endorse cortical neurons to interconnect synaptically along with receiving their previously formed synapses, such that associative memory neurons are recruited in neural pathways from the sensory cortices to the medial prefrontal cortex that encode fear-relevant anxiety.

Our data reveal that the psychological stress induces fears specific to stressful signals in battle scenes and anxiety-like behaviors by the recruitment of associative memory neurons based on the KiF1a-mediated intracellular transportation. The downregulation of KiF1a expression in the medial prefrontal cortex relieves fear memory and anxiety-like behaviors, such that KiF1a-mediated process may be one of the targets to relieve severe and persistent fear memory that may cause posttraumatic stress disorder, obsessive-compulsive disorder and generalized anxiety ^4–12^. On the other hand, the acquired fear may endorse the adaptative functions by activating the defensive behaviors in the anticipation of dangerous outcomes as well as minimizing the impact of threats ^1–3^. It is crucial to investigate the quantitative knockdown of KiF1a in order to find its borderline for maintaining moderate fear memory and relieving pathological mood ^19,20^.

The previous studies about the neural correlates for the acquired fear or fear memory have been conducted by using the fear conditioning for rodents to physically receive the electrical shocks to their feet with bell rings or contexture ^8,22–25^. As the current of electrical shocks may spread to the entire body and activate all of the neurons in the nervous system ^19,21,59,63,65,86^, the coactivity of these neurons in response to this electrical stress may cause the neural circuits to be broad interconnections. The acquired fear is also studied by the body impairment in resident/intruder paradigm ^7,21,56–61,63^. The stressful signals including somatic injury pain and battle sounds are detected by multiple sensory systems, encoded in the sensory cortices and integrated in the medial prefrontal cortex and amygdala ^21,64,65^. As body injuries are associated with pain, the cellular and molecular mechanisms underlying pain and pain-related fear may be overlapped ^18,71^. The neuronal circuits correlated to these stressful paradigms may not specifically present the features of cellular infrastructure in the acquired fear. On the other hand, compared to somatic pain-associated fear, the acquired fear more commonly occurs in observing and listening the stress episodes that cause psychological stress ^1–3,72–74^. Thus, the neural allocations and molecular profiles for the fear induced by the psychological stress should be studied ^2,7,75–78^. In addition to the amygdala ^3,74^, the visual, auditory and medial prefrontal cortices are likely involved, since these cortices encode the stressful signals and fear memories ^21,42,65,79–85^. In present study, we have looked through the basic features of neuronal infrastructures in visual, auditory and medial prefrontal cortices that are correlated to the fear induced by psychological stress. In those mice with the fear memory specific to psychological stress scenes, synapse interconnections emerge among visual, auditory and medial prefrontal cortices and among intramodal neurons. Some neurons in these cortices become able to encode the stress signals. All of these changes for the recruitment of associative memory neurons depend on KiF1a-mediated intracellular transportation. Our data provide new insights for cellular and molecular mechanisms underlying fear memory and anxiety.

The auditory cortex and the visual cortex presumably encode associative learning and memory ^81,85,118–127^. The medial prefrontal cortex is correlated to the integration of the associative memories including cognition-relevant memory and fear memory ^38,50,64,79,128–133^. The neural circuits among these cortices, the cellular infrastructure and the mechanisms related to associative memory remain unclear. Our data indicate that the emergence of psychological fear is coupled with KiF1a-mediated synapse interconnections among these cortical areas and among their intramodal neurons, which are recruited as the associative memory neurons to encode stress signals and acquired fear. These associative memory neurons to encode psychological issues in pathology functionally differ from those associative memory neurons to encode physiological signals for the cognitive activities ^87,88,91,92,134,135^. The molecular profiles to recruit associative memory neurons that encode physiological signals and the molecular profiles to recruit associative memory neurons that encode pathological signals should be addressed in order to selectively manipulate such two groups of associative memory neurons, which may benefit to maintain physiological memories and expunge pathological memories. For instance, our current study has indicated the requirements of contactin-associated protein-1 for the recruitment of associative memory neurons that encode the psychological stress and for the conversion of resilience into susceptibility to psychological stress. It is noteworthy that the molecular markers for engrams have been thought of as immediate early genes ^136–138^. These genes are nonspecific in the overly active neurons, such as excitotoxicity cells, seizure cells and pain-relevant cells ^139–145^. A partial colocalization of associative memory neurons and *cFos*-labelled neurons indicates that these two populations are not mutually represented ^65^.

In summary, the psychological stress by the formation of new synapse interconnections among cross-modal visual, auditory and medial prefrontal cortices and among intramodal neurons recruits them to be associative memory neurons that encode stress signals and fear. The associative memory neurons of encoding psychological behaviors are primary in visual and auditory cortices and secondary in the medial prefrontal cortex. The stress-induced psychological activities and cellular infrastructure changes are based upon KiF1a-mediated intracellular transportation. The coactivity of cortical neurons induced by the psychological stress towards the KiF1a-mediated processes and their featured cellular infrastructures are illustrated in Figure 8.

**Figure 8.**
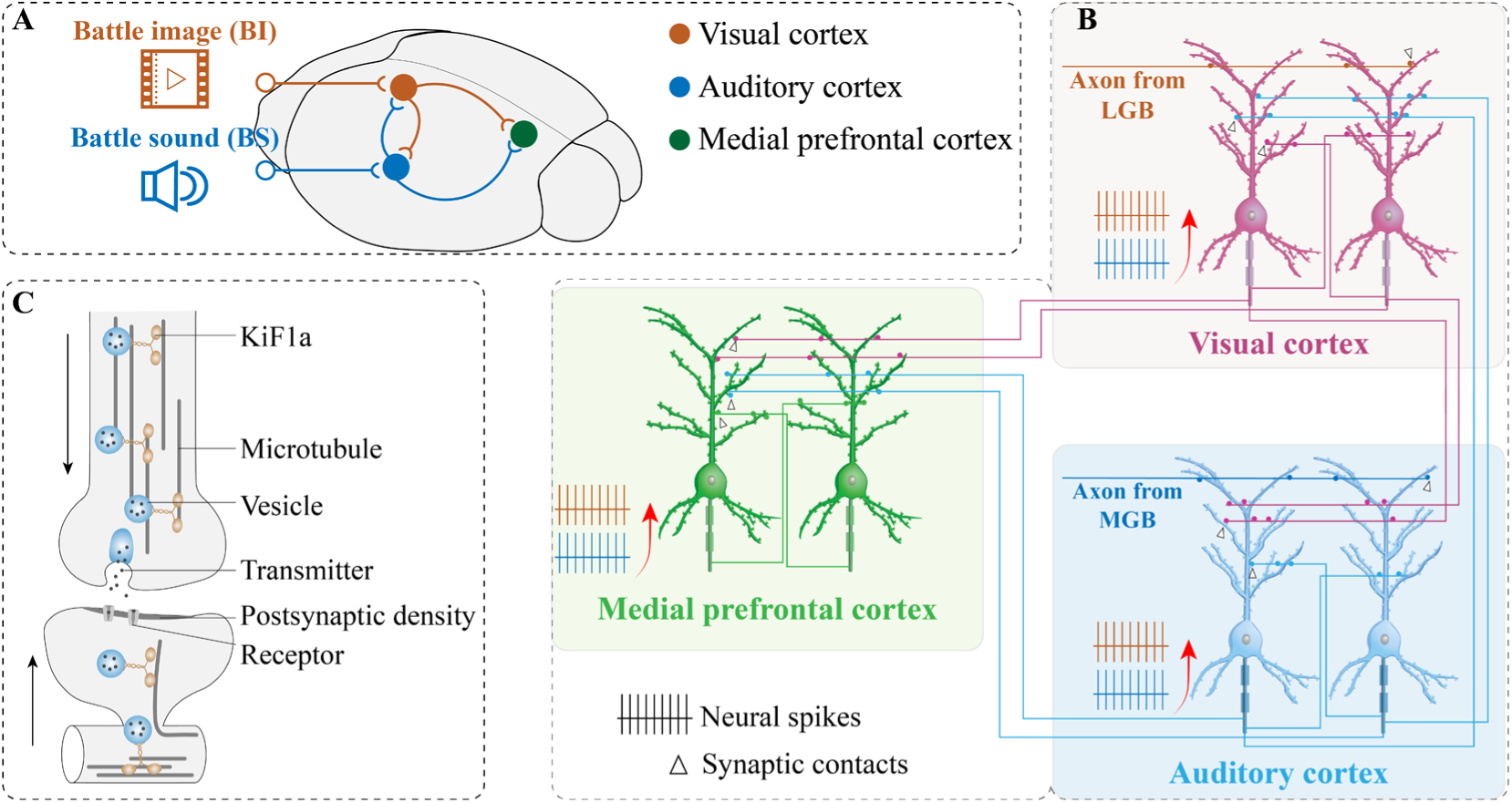
Cellular infrastructure in the auditory cortex, visual cortex and the medial prefrontal cortex correlated to the fear induced by psychological stress. **A)** The stressful signals, in which battle images (BI) are inputted to the visual cortex (VC) and battle sounds (BS) are inputted to the auditory cortex (AC), induce the synapse interconnections between auditory and visual cortical neurons as well as the convergent synapse innervations onto the neurons in the medial prefrontal cortex (mPFC) in the mice of witnessing the resident/intruder paradigm. **B)** Battle images (BI) are transmitted to the visual cortex via the axons of the lateral geniculate body (LGB). Battle sounds are transmitted to the auditory cortex via the axons of the medial geniculate body (MGB). The neurons in the visual cortex and the auditory cortex are interconnected through *en passant* synapses, or the contacts between presynaptic boutons and postsynaptic spines, after their coactivity, such that they encode battle images and battle sound, i.e., they are recruited as the primary associative memory neurons (pAMN). The neurons in the medial prefrontal cortex (mPFC) receive the convergent synapse innervations from the VC and the AC after their coactivity, such that they encode battle images and battle sound, i.e., they are recruited as the secondary associative memory neurons (sAMN). In the meantime, the intramodal interconnections via collateral axons and *en passant* synapses are strengthened among the neurons within the VC, the AC and the mPFC. After associative memory neurons are recruited, their activity strength is upregulated. **C)** KIF1A in axons and dendrites fulfills the intracellular transportations of synaptic vesicles, receptors and synapse-building elements to coordinate the axon projection, spine expansion and new synapse formation, which are essential for fear memory and anxiety-like behaviors induced by psychological stress.

## Materials and Methods

### Studies approved in mice and animal’s feeding

Experiments were conducted in accordance with the guidelines and regulations by the Administration Office of Laboratory Animals in Beijing China. All experimental protocols were approved by the Institutional Animal Care and Use Committee in Beijing China (B10831).

Two species of mice have been utilized in our experiments. C57BL/6JThy1-YFP mice (Jackson Lab, JAX stock #003782, USA) were used, whose glutamatergic neurons in the cerebral brain were labelled genetically by the yellow fluorescent protein (YFP) ^146,147^. These C57BL/6JThy1-YFP male and female mice were randomly divided into control group and observational group in face to the resident/intruder paradigm (RIP). This observational group also included the two subgroups in faces to RIP plus *KiF1a* knockdown and to RIP plus shRNA scramble control. CD1 mice above three months with more aggression were used as resident mice. The well-developed C57 mice on postnatal days (PND) 21 were selected and placed into an observation cage neighboring to the cage used for resident/intruder paradigm. These two cages were separated by a transparent and punctured wall (Figure 1A). C57 mice in observational groups were placed in the cage that was faced to a resident/intruder paradigm cage including one resident CD1 mouse and one intruder C57 mouse, in which the resident CD1 mouse attacked the intruder C57 mouse. C57 mice in control group were placed in the cage that was faced to a resident/intruder paradigm cage only including a resident CD1 mouse. That is, witness environments between observational mice and control mice are different. Observational mice watched the attacks of a resident mouse to an intruder mouse, or psychological stress in observational mice. The control mice watched resident mouse only without the stressful attack. The observer mice in observational groups by watching the attacks of a resident CD1 mouse to an intruder C57 mouse received the stressful signals including the battle scene of resident mouse attacking to intruder mouse through the visual system to the visual cortex as well as the battle sound through the auditory system to the auditory cortex, i.e. the learning of these cross-modal associated signals including battel scene and battle sound. By contrast, the control mice were exposed the cage in face to resident CD1 mouse (the scene of CD1 mouse) and to environmental tiny sound, i.e., learning of cross-modal associated signals including CD1 image and environmental sound, but not the stressful signals (battle scene and battle sound). In other words, the groups of the observational mice and control mice learnt different associated signals. These battle scenes and battle sounds were recorded for the electrophysiological experiments to test the responses and the encoding of visual, auditory and medial prefrontal cortical neurons to stressful signals.

Before facing to the resident/intruder paradigm, these observational mice and control mice were examined by the social interaction test (SIT) to identify their fear memory specific to the resident CD1 mouse and by the elevated-plus maze (EPM) test to assess their anxious state ^7,21^. The two groups of mice show no statistical differences in these tests (Figure S1). These mice were accommodated in the specific pathogen-free facility with the circadian of twelve hours for night and day plus self-help feeding. The face to resident/intruder paradigms and the behavioral tests were conducted during the daytime of this circadian.

### Behavioral study

C57 mice were fetched to the laboratory to accommodate experimenters and the training apparatus for two days and were randomly assigned into observational group and control group. The resident/intruder paradigm is introduced by the following references ^7,21,56–59,63^ and Figure 1A. In the meantime of doing resident/intruder paradigm, one observational mouse was placed in a neighboring cage with the transparent and perforated glass-plate to separate two cages (Figure 1A). The times, duration and period for observational mice to face the cage where a CD1 mouse attacked an intruder mouse in a resident/intruder paradigm were set as once a day for 30-90 seconds and twelve days in total (Figure 1A). The identical times, duration and period were given for control mice to face the cage including resident mouse. In addition, the training schedule in detail is given in supplementary table one. The protocol of this psychological stress was also modified in the stressful intensity daily during twelve-day training period, in order to reduce the habituation of observational mice to the stresses. In our studies, C57 mice in observational groups by watching the attacks of CD1 resident mouse to C57 intruder mouse received the stressful signals including the battle scene of resident mouse attacking to intruder mouse through the visual system to the visual cortex and the battle sound through the auditory system to the auditory cortex, i.e. the learning of these cross-modal associated signals including battel scene and battle sound. By contrast, the control mice were exposed the cage in face to CD1 mouse (the scene of CD1 mouse) and to environmental tiny sound, i.e., learning of cross-modal associated signals including CD1 image and environmental sound, but not the stressful signals (battle scene and battle sound). In other words, the groups of the observational mice and control mice learnt different associated signals.

The battle image in the attacks of CD1 mouse to C57 mouse in resident/intruder paradigm and the motional tracks of observational and control mice in the social interaction box were recorded by the video camera located above these open fields. The battle sound was recorded by a tiny audio recorder. That is, the complete procedure in the attacks of CD1 mice to C57 mice and their battle sound were recorded at the same time for the subsequent uses in the social interaction test and electrophysiological studies. The observational mice were sent back to their home-cage and housed individually. In control procedures, C57 mice were placed in the cage neighboring the cage for resident/intruder paradigm in the absence of intruder mouse.

The social interaction test was conducted in open field box (50 cm x 50 cm) that included small transparent and perforated box (10 cm x 10 cm) showed in Figure 1A. A CD1 mouse, mouse surrogate model or tiny audio recorder that broadcasted battle sound was placed in this small box, respectively, about six minutes to test the avoidance behaviors of observational mice and control mice as an index of the fear memories specific to CD1 mouse or battle sound. During this test, the battle sounds about one minute were given three times with one-minute intervals, which was controlled by the Bluetooth. The area within five centimeters around this small box was defined to be an interaction zone (Figure 1A, B and D). The stay duration of C57 mice within the interaction zone was named as the interaction time. The EPM consisted of two open arms (30 cm x 5 cm) crossly opposite to two closed arms (30cm × 5cm × 15.25 cm). These arms extended from a central platform (5 cm × 5 cm). The EPM was located 40 cm above the floor. The mice were placed in this central platform of the EPM in face to open arms for the test of their anxious state. The measurements of C57 mouse behavioral tests about the time in open arms and the interaction time were analyzed by a Matlab. The detail methods were described in our previous studies ^7,21,61,62,70^. The timeline for our experiments is presented in Figure 1A.

### Anterograde neural tracing

To trace the formations and changes of neural circuits in control and observational mice with fear memory induced by psychological stress, we applied adeno-associated viruses (AAV) as a vector that carried genes of encoding fluorescent proteins for an anterograde neural tracing. AAV2/8-CMV-tdTomato-WPRE, AAV2/8-CMV-EBFP2-WPRE and AAV2/8-CMV-EGFP-3xFLAG-WPRE were ordered from OBiO Technology (Shanghai, China) to track RFP-labelled, BFP-labelled and GFP-labelled axons, respectively. The titers of these viral vectors were in a range of 5-6x10^12^ genome copies/ml. All of viral vectors had been aliquoted into 200 µl EP tubes and stored at -80℃ until they were used. The locations for the microinjection of these AAVs as the original areas of neuronal axons and for the confocal scanning of cellular images as the target areas were based on a mouse brain map^148^.

In tracing axon and synapse interconnections between the visual cortex and the auditory cortex in observational and control mice, 0.3 µL AAV-CMV-tdTomato was microinjected into the visual cortex (2.50 mm posterior to the bregma, 2.50 mm lateral to the middle line and 0.35 mm depth away from the cortical surface), and 0.3 µL AAV-CMV-BFP was microinjected into the auditory cortex (2.10 mm posterior to the bregma, 3.80 mm lateral to the middle line and 0.65 mm depth away from the cortical surface) in observers three days before they witnessed the resident/intruder paradigm and in controls before they were placed neighboring the resident-included cage as well as three weeks before neural tracing was done under the confocal microscopy. The microinjections of AAVs in cerebral cortices were conducted by using glass pipettes. The microinjection quantities and durations were controlled by the microsyringe system held with three-dimensional stereotaxic apparatus (RWD Life Science, Shenzhen, China). Glass pipettes with low interior pressure were reserved at the injection sites no less than 25 minutes before withdrawing from mouse brains. In the principle, the injected AAV-CMV-tdTomato or AAV-CMV-BFP in cerebral cortices was uptaken by the somata of these cortical neurons to express red or blue fluorescent proteins, and subsequently transported toward their axonal terminals and boutons, so that the somata of neurons as the original areas and their projected axons as the target areas were tracked in an anterograde manner. The axon boutons and the contacts between the axon boutons and the dendritic spines of visual or auditory cortical neurons in target areas indicate the axons projected and the synapses formed. The new synapses plus the innate synapses convergently onto auditory and visual cortical neurons indicate that these neurons are recruited as the associative memory neurons to encode battle images and sounds. It is noteworthy that the transfection efficiency of AAVs in cerebral cortices is sufficient for neural tracing. The averaged percentage of AAV transfected neurons is similar to the averaged percentage of cFos-labelled neurons, implying AAV transfections to the active neurons^65^.

The microinjection of AAV-CMV-tdTomato and AAV-CMV-BFP in the visual cortex and the auditory cortex, respectively, was also used to track the convergent synapse innervations by their axon boutons on the dendritic spines of prefrontal cortical neurons. AAV-vectors transfected neurons in the injection areas expressed their carried-genes and fluorescent proteins. Subsequently, the fluorescent proteins produced in neuronal somata were transported over entire axons in an anterograde manner, such that their axon boutons and terminals labelled by these fluorescent proteins were detected in target areas. The contacts between the axon boutons and the dendritic spines of medial prefrontal cortical neurons were deemed as the synapse contacts. The raised contacts in experiments included the newly formed synapse contacts. Those medial prefrontal cortical neurons with the convergent synapses newly from the visual cortex and the auditory cortex were presumably associative memory neurons ^21,65,87–89,91^.

To track the axon boutons and their innervated synapses in intramodal cortices, 0.3 µL AAV-CMV-tdTomato, 0.3 µL AAV-CMV-BFP and 0.3 µL AAV-CMV-GFP were microinjected in the visual cortex (the location above), the auditory cortex (the location above) and the medial prefrontal cortex (1.70 mm anterior to the bregma, 0.30 mm lateral to the middle line and 1.30 mm depth away from the dorsal cortical surface), respectively. In the scanning by a confocal microscopy, the RFP-labelled axon boutons and their innervated contacts in the visual cortex, the BFP-labelled axon boutons and their innervated contacts in the auditory cortex, and the GFP-labelled axon boutons and their innervated contacts in the medial prefrontal cortex were thought to be intramodal in nature ^112^.

The face to a resident/intruder paradigm or the control was conducted three days after surgical operations and microinjections in order to allow C57 mice recovery from operations. In the next two weeks, the fluorescent proteins expressed in neuronal somata were transported to entire axons, axon boutons/terminals along with stress-induced axon elongations. At last, the mice were anesthetized by the intraperitoneal injections of pentobarbital sodium (60 mg/kg) and perfused through left ventricle with 25 ml 0.01M phosphate buffer solution (PBS) followed by 25 ml 4% paraformaldehyde until their bodies were rigid. The brains were quickly isolated and soaked in 4% paraformaldehyde for a fixation about 24 hours. The cerebral brains were sliced by the vibratome in a series of coronal sections with a thickness of 100 µm in the PBS. These slices were air-dried and cover-slipped with 50% glycerin in the PBS. The images about neurons, dendrites, spines and axonal boutons were taken and collected at a 60X lens for high magnification under the confocal microscope (Nikon A1R plus, Japan). The anatomic images of the cerebral brain were taken by a 4X lens for a low magnification under this confocal microscope.

In C57BL/6JThy1-YFP mice, postsynaptic neuron dendrites and spines were labelled by the YFP. The presynaptic axon boutons whose somata were infected AAV were labelled by BFP, RFP or GFP. The wavelength of the excitation laser-beam 561 nm was utilized to activate the RFP. The wavelength of the excitation laser-beam 488 was applied to activate the GFP. The wavelength of the excitation laser-beam 405 nm was used to activate the BFP. The wavelengths of the emission spectra of the BFP, GFP, YFP and RFP are 412-482 nm, 492-512 nm, 522-552 nm and 572-652 nm, respectively. Those contacts between yellow dendritic spines and red, blue or green axonal boutons with less than 0.1 µm space cleft were presumed to be the chemical synapses. The images of dendritic spines, axon boutons and synapse contacts were analyzed quantitatively by ImageJ and Imaris ^87–89,91^. Associative memory neurons were accepted by detecting two sources of boutons on the dendritic spines of YFP-labelled cortical neurons.

### Electrophysiology *in vivo*

Before the electrophysiological recording of visual, auditory or medial prefrontal cortical neurons *in vivo*, the mice in control group or observer group were anesthetized by intraperitoneal injections of pentobarbital sodium (60 mg/kg) for surgical operations after the control period or the training paradigms had been conducted. The body temperature was kept at 37 °C by a computer-controlled heating blanket. The craniotomy (1 mm in diameter) was done on the mouse skull above the left sides of the visual cortex (2.50 mm posterior to the bregma and 2.50 mm lateral to the midline), the auditory cortex (2.10 mm posterior to the bregma and 3.8 mm lateral to the midline) or the medial prefrontal cortex (1.7 mm anterior to the bregma and 0.3 mm lateral to a midline). The locations for recordings were based on mouse brain map ^148^. Electrophysiological recordings to the cortical neurons *in vivo* were conducted in the mice under light anesthetic condition with a withdrawal reflex by pinching, the eyelid blinking reflex by air-puffing and the muscle relax. The electrical discharges of unitary cortical neurons were recorded in layers IV-V of these cortices by using the glass pipettes filled with the standard solution (150 mM NaCl, 3.5 mM KCl and 5 mM HEPES). The resistance of the recording pipettes was 40-50MΩ. The electrical signals from these cortical neurons in their spontaneous spikes and their evoked spikes in response to battle sounds or battle images were recorded and acquired by an AxoClamp-2B amplifier and a Digidata 1322A, and were analyzed by the pClamp 10 system (Axon Instrument Inc. CA, USA). These spiking signals were digitized at 20 kHz and filtered by a low-pass at 5 kHz. A 100-3000 Hz band-pass filter and a second-order Savitzky-Golay filter were used to isolate spike signals. Spike frequencies were quantitatively analyzed ^87–89,91^. Battle images and battle sounds in a resident/ intruder paradigm, housing environment sounds and floral-pattern images were given for 30 seconds. The visual stimulus apparatus was an 8 cm x 16 cm LCD monitor (BOE Beijing, China) with 2400 x 1080 pixels in the resolution, the refresh rate of 60 Hz and the mean luminance of 30 cd/m^2^. The screen was positioned 12 cm from the left eye occupying ∼36° x 68° degrees of visual space. The auditory stimulus was delivered by a speaker (Huawei, Shenzhen, China) in a range of sound intensity from 60-80 dB and a range of frequencies at 220 Hz-22 kHz. The speaker was positioned 25 cm from the left ear.

Normalized spike frequency in response to one of stimuli was the number of the spike frequency in response to the stimulus in 30 seconds divided by spontaneous discharge frequency in 30 seconds before the stimulus. When the ratio reached 1.5 or above, these cortical neurons were deemed as the response to this stimulus. Associative memory neurons (AMN) were accepted by detecting a situation that those cortical neurons responded to both stimulations. The identification of associative memory neurons versus those active neurons in response to a stimulus is presented in Figure S6.

### KiF1a knockdown by shRNA carried by AAV

In the study of the role of axon transportation in the elongation of axons and the formation of new synapse innervations, KiF1a, one of the motor proteins for axon transportation ^98–100,102–104^ was knocked down in the medial prefrontal cortex by microinjecting short-hairpin RNA (shRNA) specific to KiF1a mRNA that was carried by AAV vectors (AAV-U6-sh*KiF1a*-GFP) into these cortical areas in one subgroup of observational mice. The piece of scramble sequence carried by AAV (AAV-U6-scramble-EGFP) as the control was injected in the medial prefrontal cortex in another subgroup of observational mice. These virus vectors were designed by BrainVTA (Wuhan, China). The microinjections were operated five days before training protocols for effective AAV transfection and gene knockdown (Figure S3). This approach suppresses the expression of KiF1a mRNA and protein in the medial prefrontal cortical neurons, the formation of new synapse innervations from visual and auditory cortices and the recruitment of associative memory neurons in the medial prefrontal cortex.

With the approach of KiF1a knockdown in the medial prefrontal cortex (Figure S3), the behavioral tests, AAV-mediated neural tracing and electrophysiological recording *in vivo* were done to assess the effects of the KiF1a knockdown on fear memory, synapse formation and associative memory neuron recruitment in the medial prefrontal cortex. Specifically, the encoding ability of its neurons in response to battle sounds and the battle images were analyzed and compared in two subgroups. The effects of shRNA specific to the KiF1a knockdown on new synapse formation and associative memory neuron recruitment were ensured if the numbers of new synapse contacts and associative memory neurons in KiF1a-knockdown mice were lowered significantly in comparison with scramble control mice.

### RNA extraction and quantitative real-time PCR analysis of KiF1a

The mPFC regions in the brain were isolated and harvested from control mice and observational mice twelve days after treatments. mPFC tissues in observational mice in the subgroups of KiF1a-KD and scramble control were harvested about three weeks after the bilateral injections with AAV2/9-U6-shRNA(KiF1a)-CMV-EGFP-SV40 pA or AAV2/9-U6-shRNA(scramble)-CMV-EGFP-SV40 pA (0.3 µl per side). Total RNA was isolated by using the FastPure Cell/Tissue Total RNA Isolation Kit V2 (Vazyme, Nanjing, China). The HiScript IV RT SuperMix for qPCR kit (Vazyme, Nanjing, China) was used to reversely transcribe RNAs to complementary DNAs (cDNA). qRT-PCR was performed on Applied Biosystems® QuantStudio™ 7 Flex (ABI, USA) with ChamQ Universal SYBR qPCR Master mix (Vazyme, Nanjing, China). The primer and amplicon specificity were ensured to be effective by analyzing melt curves and blasting products, DNA sequence. With *GADPH* as the internal reference, the method of 2-ΔΔCT was used to determine the relative values of mRNA expression of *KiF1a* gene. The sequences of PCR primers were 5’-CACCACTATTGTCAACCCCAAA-3’ and 5’-CCCCAATGTCCCTGTAGACCT-3’ for *KiF1a* as well as 5’-AGGTCGGTGTGAACGGATTTG-3’ and 5’-TGTAGACCATGTAGTTGAGGTCA-3’ for GADPH.

### Immunofluorescence

The mice were anesthetized by the intraperitoneal injections of urethane (1.5g/kg) and perfused via the left ventricle with ice-cold 25 ml 0.1M phosphate buffer solution (PBS, pH=7.4) followed by 25 ml 4% paraformaldehyde until their bodies were rigid. The brains of such mice were postfixed with 4% paraformaldehyde for 12 hours and then kept in the PBS. The cerebral brains were sliced by a vibratome in a series of coronal slices with 30 µm thickness of in the PBS. The slices were washed in the PBS, blocked by a buffer solution containing 5% goat serum and 0.3% Triton X-100 for 2 hours at the room temperature (22-25°C), incubated with the buffer solution that contained the primary antibody (Anti-KiF1a, Proteintech, 24841-1-AP in China) in 1% bull serum albumin (free for KiF1a staining) and 0.3% Triton X-100 at 4°C for overnight. After rinsed three times in the PBS, these slices were incubated with the secondary antibody for two hours and then, if necessary, incubated with 4′,6-diamidino-2-phenylindole for 5 minutes at room temperature. The secondary antibody for the staining was goat anti-rabbit antibody conjugated with Alexa Fluor 647 (CST, 4414). These slices were lastly rinsed three times, mounted on microscope slides, air-dried and cover-slipped with 50% glycerin in the PBS. The number of fluorescent protein-labelled neurons was counted by Image Fiji. The neurons in different channels were selected by a “threshold” tool. The selected ROI regions were added to ROI manager. An “analyze particles” (size from 1-infinity) tool was used to count the number of those neurons labelled by the KiF1a motor protein and the GFP (lower threshold level at 50, upper threshold level at 255). Images were captured under a confocal laser-scanning microscope (Nikon A1R plus, Japan). The colocalization analysis and merged images were done based on our previous studies^87–89,91^.

### Statistics

All data are presented to be arithmetic mean±SEM. The statistical analyses of our data were conducted by using GraphPad Prism 9 (a public software). The one-way ANOVA was used for the statistical comparison of the changes in behaviors, morphology and neuronal activities between those control and observer groups as well as between KiF1a-knockdown and scramble control subgroups in observers. A paired *t*-test was used to compare the activities of associative memory neurons before and after drug application, as well as the encoding ability of associative memory neurons in response to two signals. Chi-square test was used for the statistic comparison of the changes in the number and the percentage of associative memory neurons as well as active neurons in electrophysiological study among these groups. The P values equally and above 0.05 in the comparisons among the groups were set to be no statistical difference, or vice versa. One asterisk, two asterisks, three asterisks and four asterisks were presented to be P values less than 0.05, 0.01, 0.001 and 0.0001, respectively.

## Declarations

### Ethics approval and consent to participate

Experiments were done in accordance with guidelines and regulations by the Administration Office of Laboratory Animal in Beijing China. All experiment protocols were approved by Institutional Animal Care and Use Committee in this office at Beijing China (B10831).

### Consent for publication

In the publication of this manuscript, no individual’ data, such as personal details, images, videos and voice are included. All necessary consent for publication has been secured.

### Availability of data, code, and material

The datasets used and analyzed in the current study are available from the corresponding author based on the request without commercial purpose.

## Acknowledgements

This study is funded by the Natural Science Foundation of China (81971027 and U2241209) to Jin-Hui Wang.

## Authors Contributions

Jiayi Li, Yu Guo, Bingchen Chen, Yun Zhang and Lei Wang contributed to the experiments and data analyses. Jin-Hui Wang contributed to the concept, project design, funding acquisition and paper writing.

## Abbreviations

AMN: associative memory neurons
mPFC: medial prefrontal cortex
VC: visual cortex
AC: auditory cortex
BS: battle sound
BI: battle image
Ob: observer
Con: control
KiF1a: kinesin family member-1a
RIP: resident/intruder paradigm
SIT: social interaction test
EPM: elevated-plus maze
LGB: lateral geniculate body
MGB: medial geniculate body
AP: anteroposterior
D-AP5: D-(-)-2-Amino-5-Phosphonopentanoic Acid
CNQX: 6-Cyano-7-Nitroquinoxaline-2,3-Dione.

## Supplemental figures and legends

**Extended Data Figure 1.**
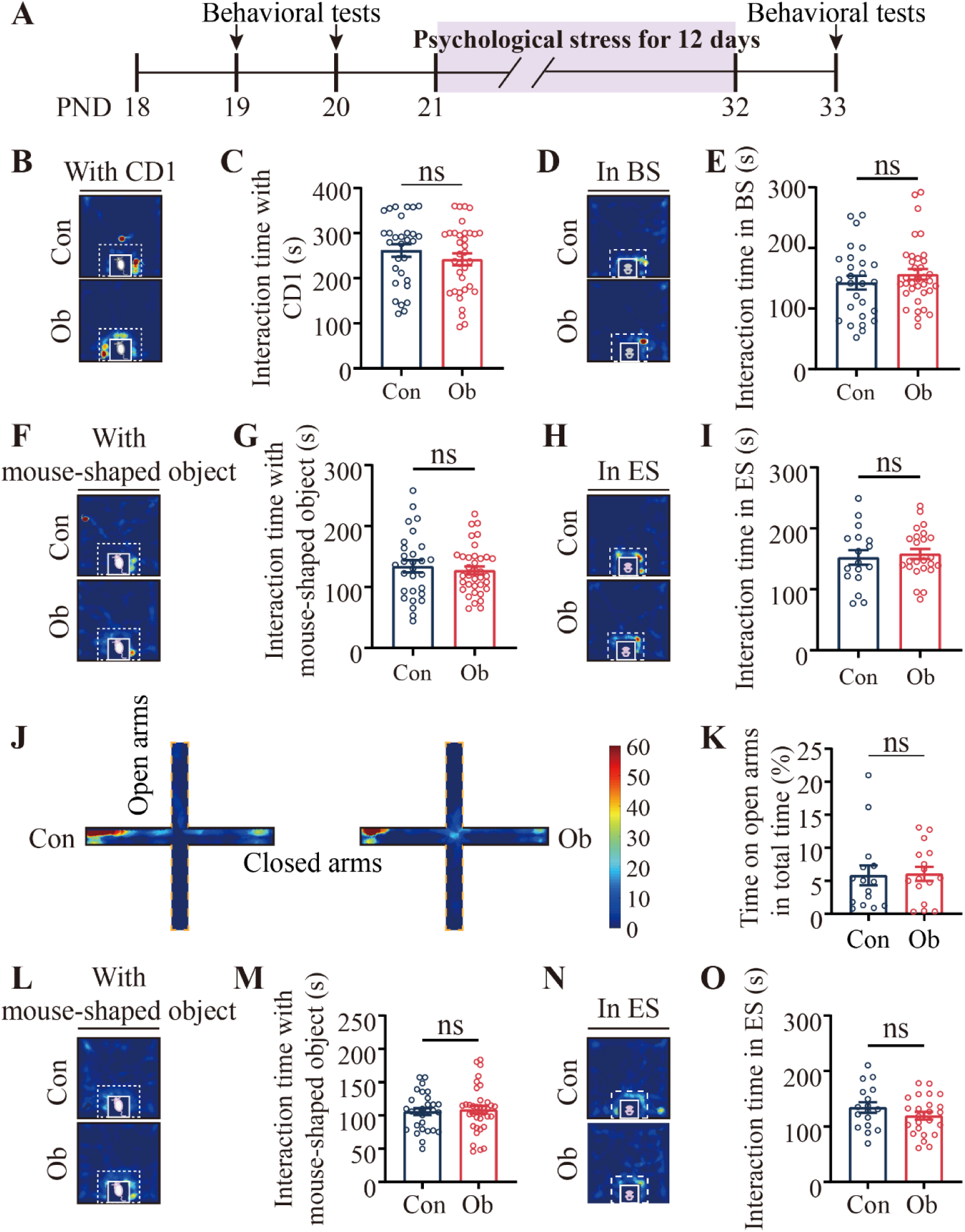
Psychological stress promotes fear memory formation specific to trauma-associated contexts in mice. **A)** Experimental timeline for mice to witness resident/intruder paradigm and to be conducted by social interaction test, elevated-plus maze test. **B)** Heat maps illustrate the social interaction test for C57 control (Con) and observational (Ob) mice in face to a CD1 resident mouse at PND 19. **C)** The bar graph with dots for each of samples show the interaction time of C57 mice with CD1 resident mouse at PND 19. **D)** Heat maps illustrate the social interaction test for C57 mice in response to battle sound at PND 19. **E)** The bar graph with dots for each of examples show the interaction time of C57 mice with the battle sound at PND 19. **F)** Heat maps illustrate the social interaction test for C57 mice in face to an artificial mouse model at PND 19. **G)** The bar graph with dots for each of samples show the interaction time of C57 mice with artificial mouse model at PND 19. **H)** Heat maps illustrate the social interaction test for C57 mice in response to housing environmental sound at PND 19. **I)** The bar graph with dots for each of examples show the interaction time of C57 mice with the housing environmental sound at PND 19. **J)** Heat maps illustrate the elevated-plus maze test for control and observational mice at PND 20. **K)** The bar graph with dots for each of samples show the time spent on open arms in the total time on the elevated-plus maze at PND 20. **L)** Heat maps illustrate the social interaction test for C57 mice in face to an artificial mouse model at PND 33. **M)** The bar graph with dots for each of samples show the interaction time of C57 mice with artificial mouse model at PND 33. **N)** Heat maps illustrate the social interaction test for C57 mice in response to housing environmental sound at PND 33. **O)** The bar graph with dots for each of examples show the interaction time of C57 mice with the housing environmental sound at PND 33. Abbreviations: PND: postnatal days; BS: battle sound; ES: housing environmental sound. ns means no statistic significant, p>0.05.

**Extended Data Figure 2.**
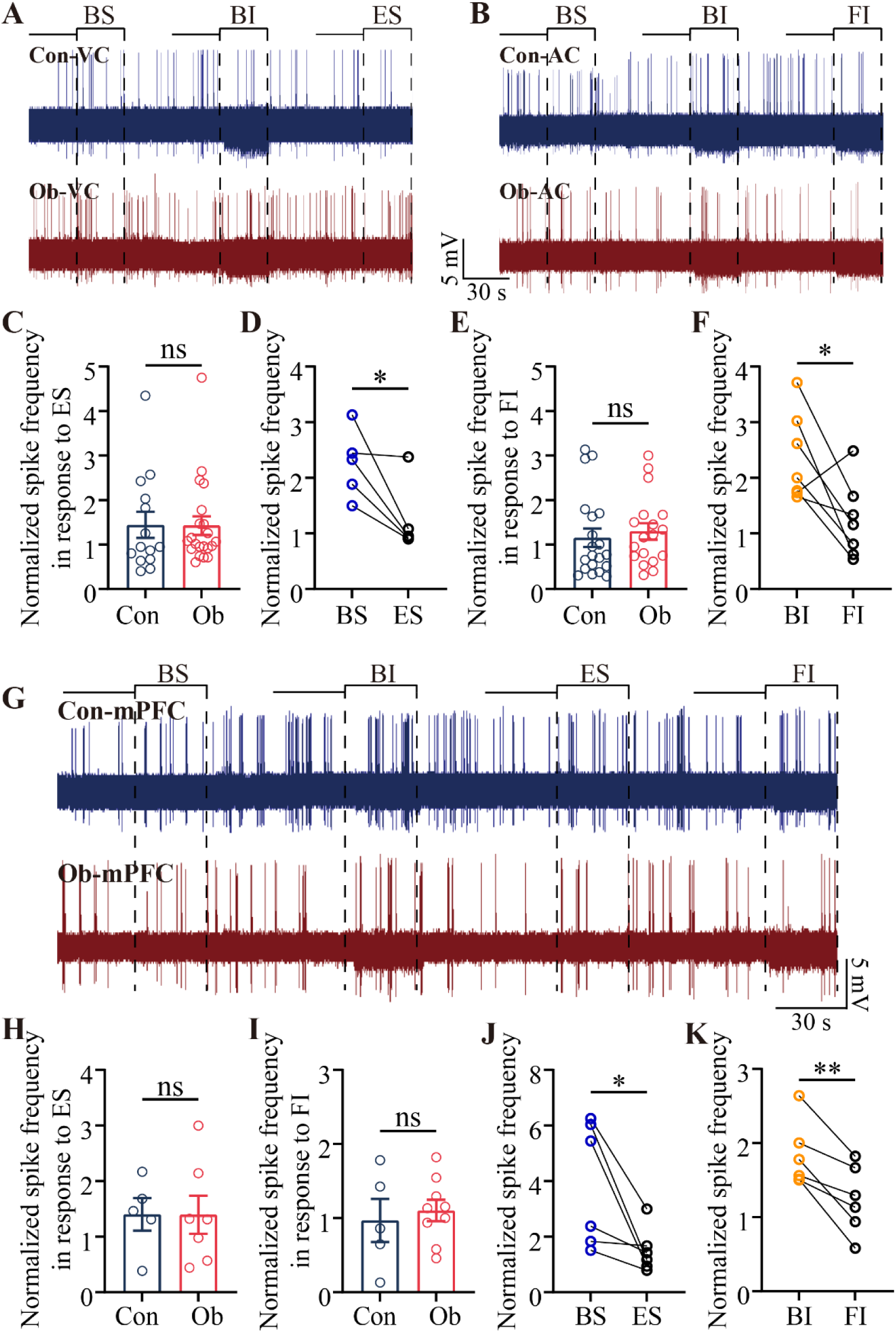
Neurons preferentially encode trauma-related information under psychological stress. **A)** The traces of spikes at VC neurons are spontaneously and evoked by BS, BI and ES from Con mouse (blue) and Ob mouse (red). **B)** The traces of spikes at AC neurons are spontaneously and evoked by BS, BI and FI from Con mouse (blue) and Ob mouse (red). **C)** Statistical comparisons about normalized spike frequencies in response to ES in VC neurons from Con and Ob mice (one-way ANOVA). **D)** Statistical comparisons about normalized spike frequencies in response to BS and ES in VC neurons (paired *t*-test). **E)** Statistical comparisons about normalized spike frequencies in response to FI in AC neurons from Con and Ob mice (one-way ANOVA). **F)** Statistical comparisons about normalized spike frequencies in response to BI and FI in AC neurons (paired *t*-test). **G)** The traces of spikes at mPFC neurons spontaneously and evoked by BS, BI, ES and FI from Con mouse (blue) and Ob mouse (red). **H)** Statistical comparisons about normalized spike frequencies in response to ES in mPFC neurons from Con and Ob mice (one-way ANOVA). **I)** Statistical comparisons about normalized spike frequencies in response to FI in mPFC neurons from Con and Ob mice (one-way ANOVA). **J)** Statistical comparisons about normalized spike frequencies in response to BS and ES in mPFC neurons (paired *t*-test) **K)** Statistical comparisons about normalized spike frequencies in response to BI and FI in mPFC neurons (paired *t*-test). Abbreviations: PND: postnatal days; BS: battle sound; ES: housing environmental sound; BI: battle image; FI: flower image. Two asterisks, p < 0.01; one asterisk, p < 0.05; ns, p >0.05.

**Extended Data Figure 3.**
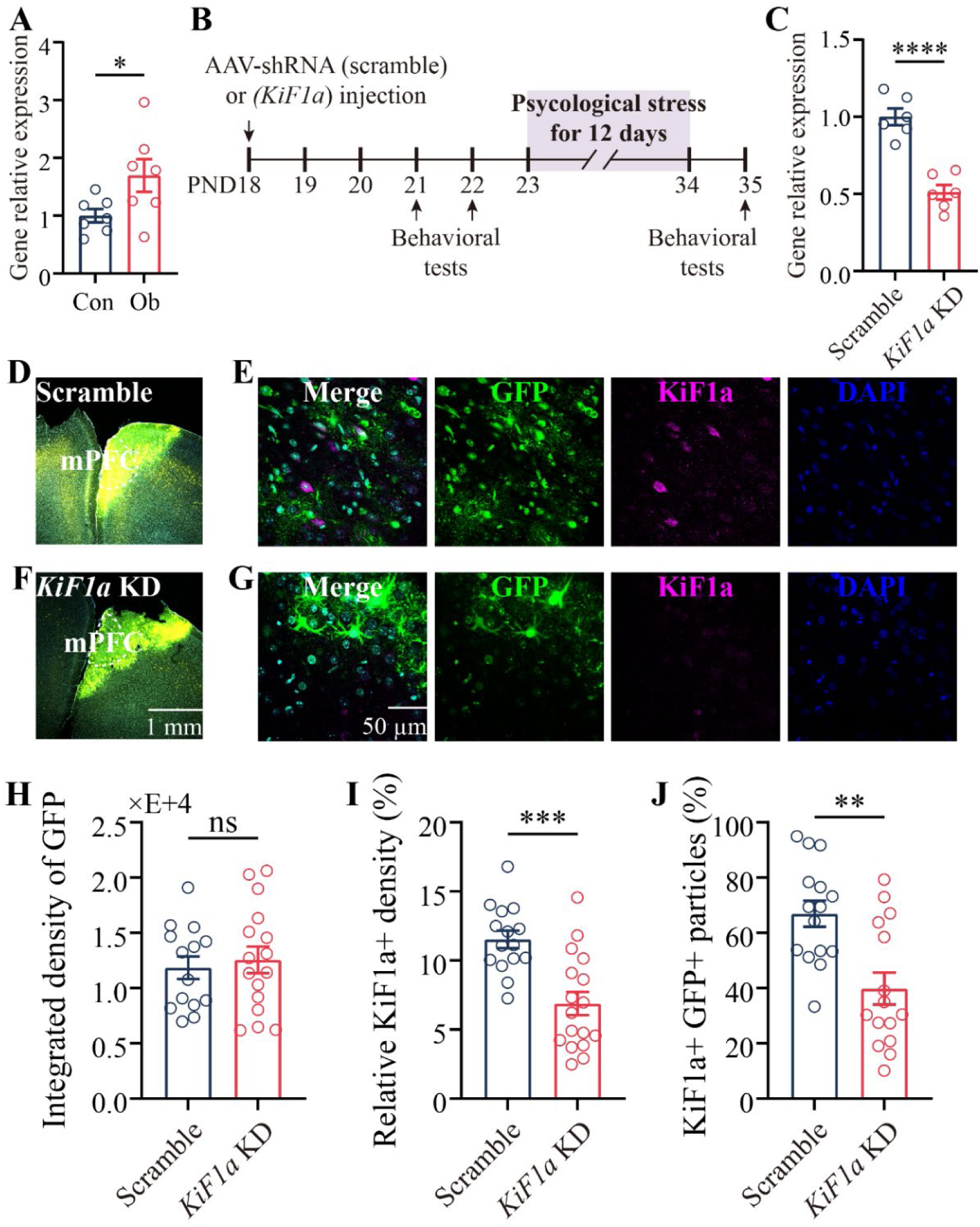
Psychological stress upregulates *KiF1a* expression in the mPFC and is suppressed by AAV-sh*KiF1a*-GFP knockdown. **A)** *KiF1a* gene relative expression in mPFC of Con and Ob mice (one-way ANOVA). **B)** Experimental timeline for RNAi performing and behavioral tests. **C)** The transcriptional level of *KiF1a* are analyzed following AAV-mediated knockdown of *KiF1a* (one-way ANOVA). **D)** Confocal image illustrates the injection sites of AAV-shRNA (scramble)-GFP. **E)** Representative confocal images of KiF1a protein level in mPFC of scramble mice. GFP labeled cells are infected with AAV-shRNA (scramble)-GFP. Magenta labeled cells are KiF1a positive cells. DAPI labeled cell nucleus. **F)** Confocal image illustrates the injection sites of AAV-shRNA (*KiF1a*)-GFP. **G)** Representative images of KiF1a protein level in mPFC of knockdown group mice. GFP labeled cells are infected with AAV-shRNA (*KiF1a*)-GFP. Magenta labeled cells are KiF1a positive cells. DAPI labeled cell nucleus. **H)** Statistic of integrated density of GFP in mPFC of scramble group mice and knockdown group mice (one-way ANOVA). **I)** Statistic of relative density of KiF1a in mPFC of scramble group mice and knockdown group mice. The data are calculated as the fluorescence intensity ratio of KiF1a to DAPI (one-way ANOVA). **J)** Statistic of the number of KiF1a and GFP double-labeled cellular particles in mPFC of scramble group mice and knockdown group mice (one-way ANOVA). The data are calculated as the ratio of KiF1a/GFP double-labeled to GFP-labeled cellular particles. Abbreviations: Con: control; Ob: observer; mPFC: medial prefrontal cortex. All data are shown as mean ± SEM. Four asterisks, p < 0.0001; three asterisks, p < 0.001; two asterisks, p < 0.01; one asterisk, p < 0.05; ns, p >0.05.

**Extended Data Figure 4.**
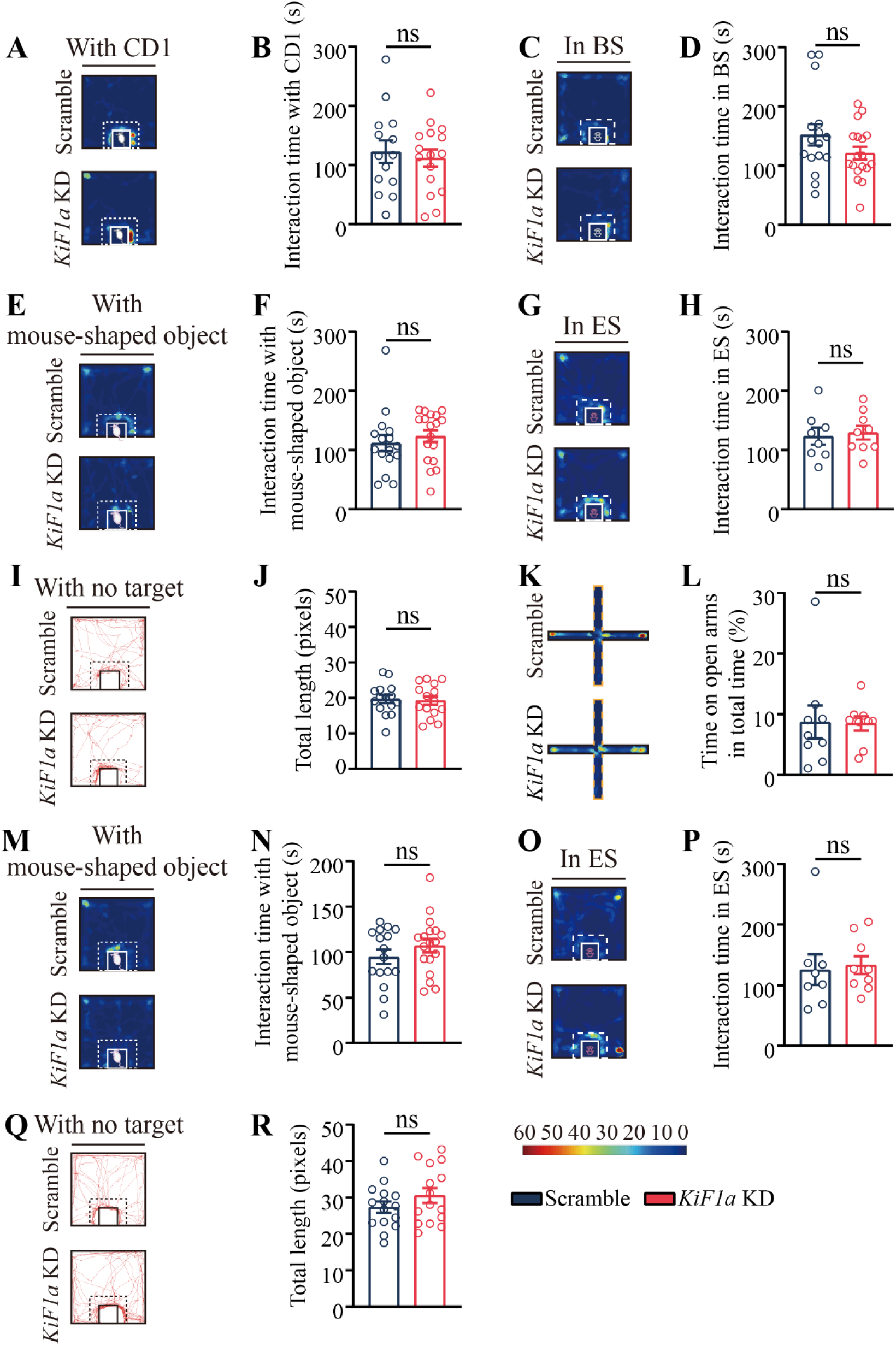
KiF1a knockdown does not affect interaction with neutral cues or locomotion. **A)** Heat maps illustrate the social interaction test for C57 scramble and *KiF1a* KD mice in face to a CD1 resident mouse at PND 21. **B)** The bar graphs with dots for each of samples show the interaction time of C57 mice with CD1 resident mice at PND 21. **C)** Heat maps illustrate the social interaction test for C57 scramble and *KiF1a* KD mice in response to battle sound at PND 21. **D)** The bar graphs with dots for each of examples show the interaction time of C57 mice with the battle sound at PND 21. **E)** Heat maps illustrate the social interaction test for C57 scramble and *KiF1a* KD mice in face to an artificial mouse model at PND 21. **F)** The bar graphs with dots for each of samples show the interaction time of C57 mice with artificial mouse model at PND 21. **G)** Heat maps illustrate the social interaction test for C57 mice in response to housing environmental sound at PND 21. **H)** The bar graphs with dots for each of examples show the interaction time of C57 mice with the housing environmental sound at PND 21. **I)** Locomotion traces of PND 21 scramble and *KiF1a* KD mice in the social interaction test. **J)** The bar graphs with dots for each of samples show the total movement trajectory length of C57 mice at PND 21. **K)** Heat maps illustrate the elevated-plus maze test for scramble and *KiF1a* KD mice at PND 22. **L)** The bar graphs with dots for each of samples show the time spent on open arms in the total time on the elevated-plus maze at PND 22. **M)** Heat maps illustrate the social interaction test for C57 scramble and *KiF1a* KD mice in face to an artificial mouse model at PND 35. **N)** The bar graphs with dots for each of samples show the interaction time of C57 mice with artificial mouse model at PND 35. **O)** Heat maps illustrate the social interaction test for C57 mice in response to housing environmental sound at PND 35. **P)** The bar graphs with dots for each of examples show the interaction time of C57 mice with the housing environmental sound at PND 35. **Q)** Locomotion traces of PND 35 scramble and *KiF1a* KD mice in the social interaction test. **R)** The bar graphs with dots for each of samples show the total movement trajectory length of C57 mice at PND 35. All data are shown as mean ± SEM and analyzed by one-way ANOVA. Blue symbols show the data of scramble group mice. Red symbols show the data of *KiF1a* KD group mice. Abbreviations: PND: postnatal days; BS: battle sound; ES: housing environmental sound. ns, p >0.05.

**Extended Data Figure 5:**
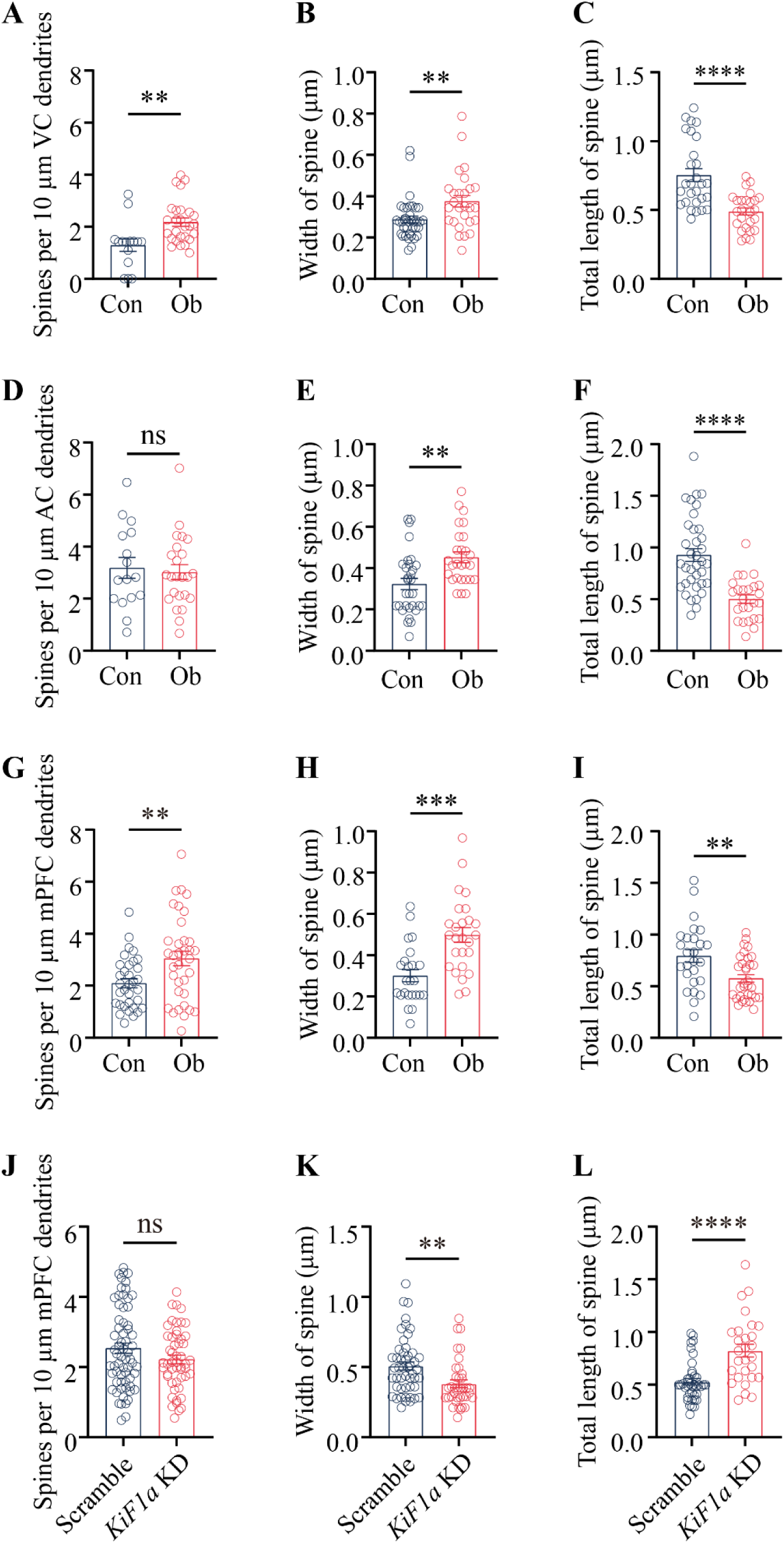
Spine morphology changes in control, Ob, scramble, and *KiF1a* RNAi groups. **A)** Statistic on the number of spines in VC of Con and Ob mice. **B)** Statistic on the width of spine in VC of Con and Ob mice. **C)** Statistic on the total length of spine in VC of Con and Ob mice. **D)** Statistic on the number of spines in AC of Con and Ob mice. **E)** Statistic on the width of spine in AC of Con and Ob mice. **F)** Statistic on the total length of spine in AC of Con and Ob mice. **G)** Statistic on the number of spines in mPFC of Con and Ob. **H)** Statistic on the width of spine in mPFC of Con and Ob mice. **I)** Statistic on the total length of spine in mPFC of Con and Ob mice. **J)** Statistic of the number on spines in mPFC of scramble and *KiF1a* KD mice. **K)** Statistic on the width of spine in mPFC of scramble and *KiF1a* KD mice. **L)** Statistic on the length of spine in mPFC of scramble and *KiF1a* KD mice. Abbreviations: Con: control; Ob: observer; VC: visual cortex; AC: auditory cortex; KD: knock down. Data are shown as mean ± SEM and analyzed by one-way ANOVA. Four asterisks, p < 0.0001; three asterisks, p < 0.001; two asterisks, p < 0.01; ns, p >0.05.

**Extended Data Figure 6:**
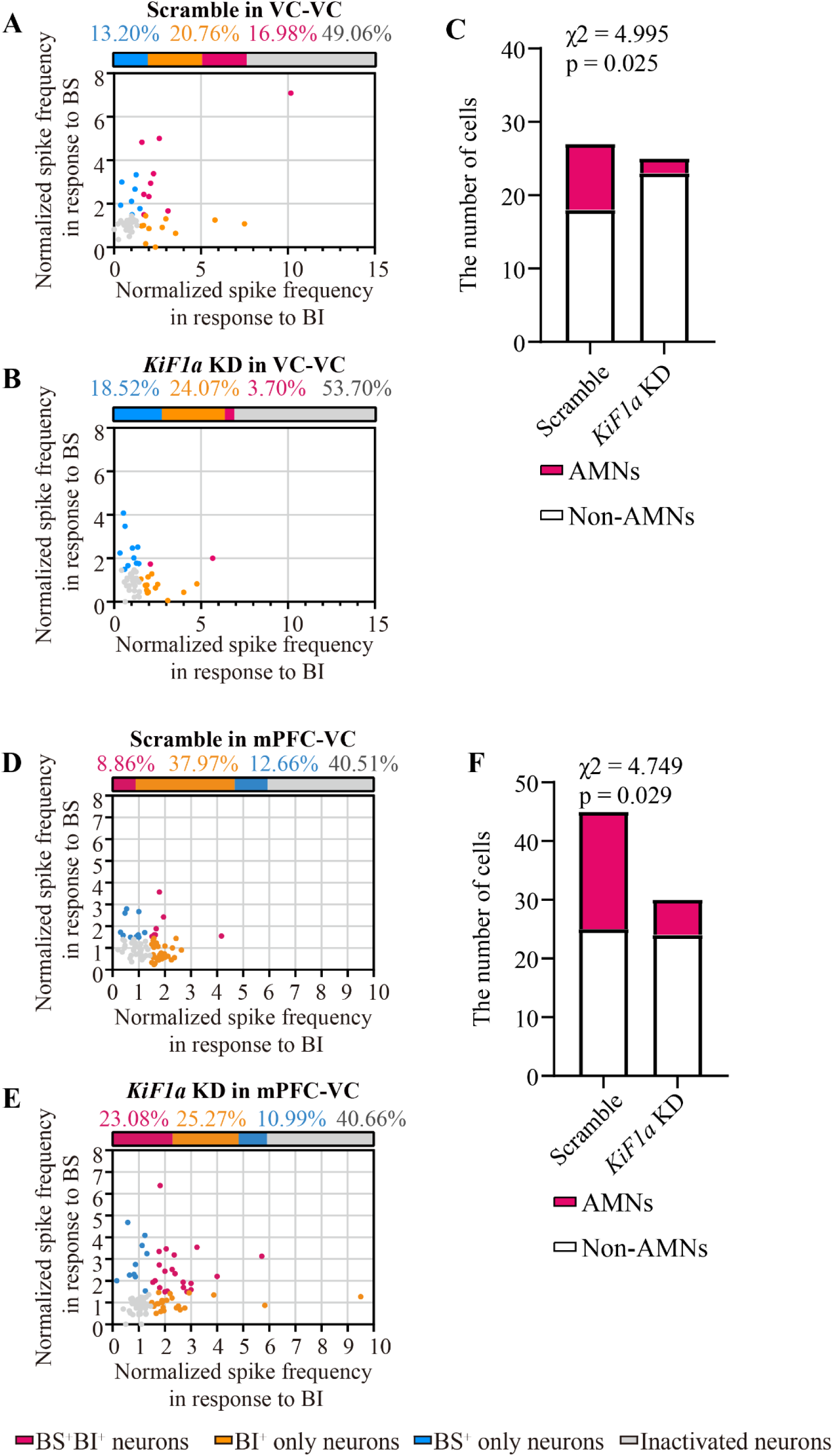
*KiF1a* knock down in VC or mPFC reduces formation of AMNs in VC. **A)** The scatter plot illustrates the encoding capacity of BS and BI information by individual neurons in the VC region following shRNA(scramble) injection into the same area during *in vivo* electrophysiological recordings. **B)** The scatter plot illustrates the encoding capacity of BS and BI information by individual neurons in the VC region following sh*KiF1a* injection into the same area. **C)** Statistical comparations about the numbers of associative memory neurons (AMNs, red in bar) that encode both BI and BS as well as of activated neurons that encode either BI or BS (non-AMNs, white in bar) in the visual cortex from Scramble and *KiF1a* KD mice (AAV-shRNA is injected into VC, chi-square test) **D)** The scatter plot illustrates the encoding capacity of BS and BI information by individual neurons in the VC region following shRNA(scramble) injection into the mPFC brain area during *in vivo* electrophysiological recordings. **E)** The scatter plot illustrates the encoding capacity of BS and BI information by individual neurons in the VC region following sh*KiF1a* injection into the mPFC brain area. **F)** Statistical comparations about the numbers of associative memory neurons (AMNs, red in bar) that encode both BI and BS as well as of activated neurons that encode either BI or BS (non-AMNs, white in bar) in the visual cortex from Scramble and *KiF1a* KD mice (AAV-shRNA is injected into mPFC, chi-square test) Each dot on the scatter plot represents a cell recorded using in vivo electrophysiological techniques. The blue dots represent neurons selectively responsive to battle sound. The yellow dots represent neurons selectively responsive to battle image. The red dots represent neurons respond to both battle sound and battle image. The grey dots indicate inactivated neurons that respond to neither battle sound nor battle image. Abbreviations: AMNs: associative memory neurons; BS: battle sound; BI: battle image.

---

**Table 1.** Schedule for psychological stress in C57 mice in face to resident/intruder paradigm.

| Days<br>Weeks | One | Two | Three | Four | Five | Six | Seven |
| --- | --- | --- | --- | --- | --- | --- | --- |
| One | ORIP (30 s)<br>Post-LSS (30 s) | ORIP (60 s),<br>Post-LSS (30 s) | ORIP (60 s) | ORIP (60 s),<br>Post-LSS (60 s) | ORIP (60 s),<br>Post-LSS (60 s) | Pre-LSS (30 s),<br>ORIP (60 s)<br>Post-LSS (60 s) | ORIP (60 s)<br>Post-LSS (90 s) |
| Two | ORIP (60 s)<br>Post-LSS (120 s) | ORIP (90 s),<br>Post-LSS (60 s) | Pre-LSS (60 s),<br>ORIP (60 s),<br>Post-LSS (120 s) | Pre-LSS (60 s)<br>ORIP (60 s),<br>Post-LSS (120 s) | Pre-LSS (60 s)<br>ORIP (90 s)<br>Post-LSS (120 s), |  |  |

**Definition:** The fear learning is to observe the attacks of a CD1 mouse to an intruder mouse in the resident/intruder paradigm about 30-60 seconds. The pre-learning of stressful situations is for observational mouse with intruder mouse to observe CD1 mouse about 30-60 seconds, The post-learning of stressful situations is for observational mouse and intruder mouse to communicate each other about 30-120 seconds. To enhance psychological stress behavioral training, we incorporated two modules: Ob/Intruder co-observation of CD1 mice (30-60 s, pre-LSS) and Ob-injured intruder mouse interaction (30-120 s, post-LSS).
**Abbreviations:** ORIP, observation to resident/intruder paradigm; Pre-LSS, the pre-learning of stressful situation. Post-LSS, the post-learning of stressful situations

